# The social cerebellum: A large-scale investigation of functional and structural specificity and connectivity

**DOI:** 10.1101/2021.02.15.431044

**Authors:** Athanasia Metoki, Yin Wang, Ingrid R. Olson

**Affiliations:** Department of Psychology, Temple University, Philadelphia, USA; State Key Laboratory of Cognitive Neuroscience and Learning, and IDG/McGovern Institute for Brain Research, Beijing Normal University, Beijing, China

**Author notes:** Address correspondence to: Dr. Athanasia Metoki, Dr. Ingrid Olson.

**Keywords:** cerebellum, theory of mind, effective connectivity, structural connectivity, human connectome project

## Abstract

The cerebellum has been traditionally disregarded in relation to non-motor functions, but recent findings indicate it may be involved in language, affective processing, and social functions. Mentalizing is the ability to infer mental states of others and this skill relies on a distributed network of brain regions. Here, we leveraged large-scale multimodal neuroimaging data to elucidate the structural and functional role of the cerebellum in mentalizing. We used functional activations to determine whether the cerebellum has a domain-general or domain-specific functional role, and effective connectivity and probabilistic tractography to map the cerebello-cerebral mentalizing network. We found that the cerebellum is organized in a domain-specific way and that there is a left cerebellar effective and structural lateralization, with more and stronger effective connections from the left cerebellar hemisphere to the right cerebral mentalizing areas, and greater cerebello-thalamo-cortical (CTC) and cortico-ponto-cerebellar (CPC) streamline counts from and to the left cerebellum. Our study provides novel insights to the network organization of the cerebellum, an overlooked brain structure, and mentalizing, one of humans’ most essential abilities to navigate the social world.

## Introduction

The cerebellum has traditionally been considered a motor brain area, involved in the coordination of voluntary movements, gait, posture, and speech (Glickstein 1992, 1993; Brodal and Bjaalie 1997; Fine et al. 2002; Ito 2002). In more recent years there has been a newfound interest in non-motor functions of the cerebellum (Leiner et al. 1986; Schmahmann and Pandya 1989, 1997; Schmahmann 1991, 1998; Middleton and Strick 1994). This work has clinical grounding in findings showing that cerebellar damage from stroke or injury could at times cause the “Cerebellar Cognitive Affective Syndrome” which includes language deficits such as mutism, dysprosodia, and anagrammatism, problems with executive functioning, as well as personality changes, blunting of affect, and inappropriate social behavior (Schmahmann and Sherman 1998). More strikingly, resection of the most common type of pediatric brain tumors which are found in the cerebellum, at times results in dramatic and long-lasting changes in cognition as well as personality, affect, and overall social behavior (Albazron et al. 2019).

Buckner and colleagues used resting-state analyses at the group level and showed, that the cerebellum has a distinct mapping of all the large-scale networks found in the cerebrum with each resting-state network represented thrice, specifically in lobules VI-Crus I, Crus II-VIIb, and lobule IX (Buckner et al. 2011). Guell, Gabrieli, et al. (2018) confirmed and extended Buckner’s demonstration using task-based fMRI data from the Human Connectome Project dataset. Lastly, (King et al. 2019) used a multi-domain task battery and derived a comprehensive functional parcellation of the cerebellar cortex. We can distill these findings into a few take-homes: (1) motor functions are localized to the anterior lobe and inferior posterior lobe; cognitive functions to the superior posterior lobe; emotion/affect potentially to the vermis; (2) there are finer-scale functional subregions within each lobe; and (3) the functional subregions do not neatly follow sulcal/gyral boundaries.

Furthermore, research confirmed the existence of a cerebello-cerebral structural network. Early animal studies have shown that the white matter connections of this structural network are predominately contralateral and form closed loops (Kelly and Strick 2003). The cerebellum projects to parts of the contralateral cerebral cortex via the thalamus, and then receives input from the same cerebral areas which project back to the cerebellum via the pons, bypassing the thalamus. More recent animal studies confirmed this finding but also found ipsilateral cerebello-cerebral connections (Suzuki et al. 2012). These findings were confirmed by diffusion imaging research in humans which showed that contralateral connections comprise ∼70-80% of cerebello-cerebral white matter connectivity with the remainder being ipsilateral (Krienen and Buckner 2009; Salmi et al. 2010; Sokolov et al. 2014).

What is the best way to describe the role of the cerebellum in social cognition? Van Overwalle and colleagues performed meta-analyses on fMRI data and showed that portions of the cerebellum are consistently activated in mentalizing tasks (Van Overwalle, Baetens, et al. 2015; Van Overwalle, D’aes, et al. 2015). Mentalizing is the ability to attribute mental states to others, and to interpret their intentions, perspectives, and beliefs (Frith and Frith 2006; Blakemore 2008; Mar 2011; Schurz et al. 2014) and is an essential social skill. This finding is interesting because there has been a concerted effort in social neuroscience over the last twenty years to identify brain areas involved in mentalizing. Researchers have found that the “mentalizing network” is comprised of portions of the temporoparietal junction, precuneus, amygdala, anterior temporal lobe, occipital gyrus, fusiform gyrus, inferior frontal gyrus, and medial prefrontal cortex (Uddin et al. 2007; Lahnakoski et al. 2012; Bzdok et al. 2013, 2015; Yang et al. 2015; Hartwright et al. 2016) and these regions are interconnected by a system of white matter pathways (Wang et al. 2020). However, with the exception of work by Van Overwalle and colleagues, the cerebellum has been largely ignored by researchers interested in mentalizing and it is not included in the classic mentalizing network.

In sum, the lesion findings are compelling, but they do little to illuminate what part of the cerebellum is involved in social and affective behavior. The neuroimaging findings are scarce, with few in-depth studies looking at large scale networks so we lack an understanding of how the cerebellum interacts with social and affective regions in the cerebrum. In the present study, we use a large multi-modal fMRI dataset to investigate the cerebellar foundations of mentalizing. Using clustering methods, effective connectivity, and probabilistic tractography we show that the cerebellum is divided into distinct functional parcels, strongly connected to individual tasks. Furthermore, we show that there are mentalizing cerebellar areas effectively connected specifically to regions of the cerebral cortex that have a known role in mentalizing. Last, some – but not all – portions of this functional network are supported by an underlying white matter network.

## Materials and Methods

### Dataset and Participants

All data used in this study are part of the Human Connectome Project (HCP) dataset, specifically the WU-Minn HCP Consortium S900 Release (WU-Minn HCP Consortium 2015). This dataset is publicly available, accessible at https://www.humanconnectome.org. Only subjects that completed all imaging sessions of interest (T1/T2, task fMRI (tfMRI), and diffusion MRI (dMRI)) were included in this study. To reduce variance in structural organization (McKay et al. 2017) we restricted our population to only right-handed subjects using the Edinburgh Handedness questionnaire (Oldfield 1971), which resulted in 679 healthy young adults. Additionally, seven subjects were excluded due to lack of enough robust signal in all bilateral regions of interest (ROIs) in the mentalizing localizer task, and one more subject for missing all the explanatory variable files with timing information necessary for psychophysiological interaction (PPI) analyses, thus leading to a final sample of 671 healthy young adults (377 females, *M* = 28.8, *SD* = 3.7). Unless otherwise stated, all significant results reported in this study were corrected for multiple comparisons using the false discovery rate (FDR; Benjamini and Hochberg 1995).

### Overview of HCP protocol

Due to the complexity of the HCP data acquisition and preprocessing pipelines, listing all scanning protocols and data analysis procedures in detail is beyond the scope of this article. Full detailed description of all protocols and procedures can be found elsewhere (Van Essen et al. 2012; Barch et al. 2013; Glasser et al. 2013; Smith et al. 2013). Briefly, the HCP protocol includes acquisition of structural MRI, resting-state and task-state fMRI, diffusion MRI, and extensive behavioral testing. Task-state fMRI encompasses seven major domains: 1) social cognition (mentalizing); 2) motor (visual, motion, somatosensory, and motor systems); 3) gambling; 4) working memory/cognitive control systems and category specific representations; 5) language processing (semantic and phonological processing); 6) relational processing; and 7) emotion processing (Barch et al. 2013). The imaging data used in this article are the “minimally preprocessed” included in the WU-Minn HCP Consortium S900 Release (WU-Minn HCP Consortium 2015). Details of imaging protocols, preprocessing pipelines, and in-scanner task protocols can be found in the Supplementary Material.

As the main task of interest, participants in the mentalizing task viewed 20s video clips of geometrical shapes that either interacted with each other in a socially meaningful way or moved purposelessly on the screen. The video clips of this Heider and Simmel-type task were developed by Castelli et al. (2000) and Wheatley et al. (2007) and have been validated as a measure of mentalizing given evidence that they generate task related activation in brain regions associated with mentalizing with reliable results across subjects (Castelli et al. 2000, 2002; Wheatley et al. 2007; White et al. 2011; Barch et al. 2013). Both the emotion and the mentalizing task require some degree of mentalizing. The mentalizing task asks participants to implement mental state attributions when watching the geometrical shapes interact, hence the mentalizing is intentional. On the other hand, the emotion task has a far smaller degree of mentalizing as participants only implicitly attribute mental states to the faces they view (Van Overwalle and Vandekerckhove 2013; Kliemann and Adolphs 2018).

### Regions of Interest (ROIs)

We used three sets of ROIs for our analyses: one set in the cerebrum and two in the cerebellum. The cerebral brain regions were drawn from prior work on mentalizing (Schurz et al. 2014; Wang et al. 2020) thus the ROIs included the anterior temporal lobe (ATL; Olson et al. 2007; Ross and Olson 2010; Wang et al. 2017), dorsomedial prefrontal cortex (DMPFC; Van Overwalle 2009; Van Overwalle and Baetens 2009; Van Overwalle and Vandekerckhove 2013; Schurz and Perner 2015), precuneus (PreC; Cavanna and Trimble 2006; Uddin et al. 2007; Schiller et al. 2009; Schurz et al. 2014; Peer et al. 2015), temporoparietal junction (TPJ; Van Overwalle 2009; Koster-Hale and Saxe 2013), and ventromedial prefrontal cortex (VMPFC; (Molenberghs et al. 2016; Koster-Hale et al. 2017). Although other brain areas support the mentalizing network, we focused on brain areas involved in basic, core social processes. To ensure sensitivity within each individual, we defined individual-specific ROIs using the social cognition task as a mentalizing ROI localizer. To precisely localize each mentalizing ROI, the social cognition task was processed on the “grayordinate-based” space (cortical surface vertices and subcortical voxels) using the MSM-All registration (Robinson et al. 2014). We used the Connectome Workbench software (Marcus et al. 2011) to manually extract the vertices (which were later transformed to MNI coordinates) of the peak activations of the bilateral ten predefined ROIs (as well as their magnitudes) from the contrast “ToM > random” (socially meaningful interaction of the shapes > random movement of the shapes) for each individual separately.

These individual-specific cluster peak coordinates were used as input (spheres, 6 mm radius) for subsequent seed-based brain connectivity analyses at the individual level (PPI and probabilistic tractography). It is important to note that, based on the meta-analysis by Schurz et al. (2014), which showed that activation from the social or intentional interactions > physical movements contrast spans broadly on the TPJ, the TPJ was defined as a broad brain area encompassing the inferior parietal lobule (IPL, i.e. perspective taking) and posterior superior temporal sulcus (pSTS, i.e. biological motion perception). In the event that the TPJ had multiple clusters (falling either into what is traditionally considered to be the TPJ, or IPL or pSTS), we selected the strongest cluster to avoid excessive inter-subject inconsistency.

For the ROIs in the cerebellum we used a different approach. The cerebellum has uniform cortical cytoarchitecture and although several studies have attempted to functionally map the cerebellar cortex (Krienen and Buckner 2009; Buckner et al. 2011; Diedrichsen and Zotow 2015; Riedel et al. 2015; Guell, Gabrieli, et al. 2018; Guell, Schmahmann, et al. 2018; Marek et al. 2018; King et al. 2019), there is no consensus about functional boundaries within the cerebellum. Hence, it was impossible to employ the same approach as we did with the predefined cerebral ROIs. Instead we used a data-driven approach. Following the method used by Guell, Gabrieli, and Schmahmann (2018), we transformed individual level 2 cope files (results of within-subject fixed-effects grayordinate-based analyses which generate output files that index mean effects for an individual subject averaged across the two scan runs for a task) into Cohen’s d group maps by first transforming the grayordinate .dscalar.nii files to NIfTI. We then used FSL commands fslselectvols to extract the contrast of interest “ToM > random” for each individual, and fslmerge, fsmaths -Tmean, -Tstd, and -div to merge the individual contrast images, extract the mean, and the standard deviation, and divide the two, ultimately getting group Cohen’s d maps for the contrasts “ToM > random” (mentalizing), “faces > shapes” (emotion), “story > math” (language), “2-back > 0-back” (working memory), “reward > punishment” (gambling), “relational > match” (relational processing), and “average” (motor) based on our 671 subjects. The HCP S900 Release provides level 3 group z-maps, but Cohen’s d maps made it possible to observe the effect size of each task contrast rather than the significance of the BOLD signal change. A sample of 671 subjects ensures that a d value higher than 0.5 (Cohen 1988) will be statistically significant even after correction for multiple comparisons (d = z/sqrt(n), *d* > 0.5 we have *z* > 12.95 for *N* = 671; analysis of 17,853 cerebellar voxels would require *P* < 0.000028 after Bonferroni correction, and *P* < 0.000028 is equivalent to *z* > 4.026). Accordingly, we used the Cohen’s d maps and a threshold of 0.5 to extract clusters of activation for each task and local maxima within each cluster. After using a whole cerebellar mask to retain only the clusters and local maxima within the cerebellum, clusters smaller than 100 mm^3^ were further removed in order to omit very small clusters that were considered to be non-informative and would make a comprehensive description of the results too extensive. The coordinates of the remainder local maxima from the “ToM > random” (mentalizing) contrast within the cerebellum were used to create group cerebellar ROIs (spheres, 6 mm radius). Eleven mentalizing ROIs were created in the left cerebellar hemisphere and seven ROIs were created in the right cerebellar hemisphere. The same method was used to extract the cerebellar motor ROIs which were used in our effective and structural connectivity control analyses. Seven motor ROIs were created in the left cerebellar hemisphere and seven were created in the right cerebellar hemisphere.

### Cluster Overlap and Euclidean distances

Given that the HCP dataset uses FNIRT registration to the MNI template, we calculated the percentage of overlap of each cerebellar cluster, in reference to the cerebellar lobules, by using Diedrichsen’s FNIRT MNI maximum probability map (Diedrichsen et al. 2009). It has been shown though that functional parcellation of the cerebellum does not match its anatomical parcellation into lobules (King et al. 2019). Because of this, we created an atlas of cerebellar lobes (anterior, posterior, flocculonodular, and vermis) by combining the lobules from Diedrichsen’s FNIRT MNI maximum probability map (Diedrichsen et al. 2009) that belong in each lobe and used these maps for our analyses. We used FSL’s atlasq tool to determine the percentage of overlap of each cerebellar cluster to the cerebellar lobes, hence determining the primary location of each cluster. The Sørensen–Dice coefficient, which is a statistic measuring the similarity of two samples (Dice 1945; Sørensen 1948), was then used to calculate the percentage of overlap between the functional clusters generated from all tasks and determine their similarity, and Euclidean distances were calculated to estimate the distances of local maxima within and between clusters. At the individual level, we thresholded *Z*-scored *β*-weights of each subject’s activation map for each task contrast to > 0 to retain only increased activation during the tasks and then ran Wilcoxon signed-rank tests between each task pair to examine whether there was a statistical difference between them.

### Psychophysiological Interaction (PPI) Analyses

PPI analyses were used to understand effective connectivity – a way to capture stimulus-driven patterns of directional influence among neural areas (Friston et al. 1997) – by identifying brain regions whose activity depends on an interaction between psychological context (the task) and physiological state (the time course of brain activity) of the seed region (Gerchen, Bernal-Casas, & Kirsch, 2014; O’Reilly, Woolrich, Behrens, Smith, & Johansen-Berg, 2012; Smith, Gseir, Speer, & Delgado, 2016). We built a generalized PPI model (McLaren et al. 2012) using a non-deconvolution method (Di and Biswal 2017) in FSL for each cerebellar and cerebral ROI. Our model had five separate regressors: two psychological regressors of task events (mental interaction and random interaction), one physiological regressor of the time series of the seed ROIs and two corresponding interaction regressors (task events × seed ROI’s time series). To estimate the effective connectivity, we used the contrast between the interaction regressors (PPI mental > PPI random). *Z*-scored *β*-weights were extracted for each pair of cerebellar to contralateral cerebral ROIs, which resulted in an 11 x 5 matrix for the left cerebellar-right cerebral hemispheres and a 7 x 5 matrix for the right cerebellar-left cerebral hemispheres for each individual. We applied symmetrization to the matrices by averaging the *Z*-scored *β*-weights of each pair of cerebro-cerebellar and cerebello-cerebral ROIs (Tompson et al. 2020). At the group level of the effective connectivity, one-sample t-tests were performed across individuals at each pair of ROIs to detect any significant effective connections. What we expected to learn from this analysis is which cerebellar mentalizing regions present significant coupling with contralateral cerebral mentalizing regions due to mental state attribution (mentalizing task). The same effective connectivity analyses were performed between the mentalizing cerebral ROIs and motor (control) cerebellar ROIs.

### Diffusion Analyses

Probabilistic tractography analyses were performed using FSL’s probtrackx2 (probabilistic tracking with crossing fibres) (Behrens et al. 2003, 2007) in each subject’s native space and then the results were transformed to MNI standard space. An ROI-to-ROI approach was used with cerebellar and contralateral cerebral ROIs used as seeds and targets to reconstruct each subject’s cerebello-cerebral white matter connections. Fiber tracking was initialized in both directions separately (from seed to target and vice versa) and 5,000 streamlines were drawn from each voxel in each ROI. Tractographies were performed to delineate the cerebello-thalamo-cortical (CTC) (Middleton and Strick 1997; Schmahmann and Pandya 1997; Palesi et al. 2017) and cortico-ponto-cerebellar (CPC) (Ramnani 2006; Palesi et al. 2017) between left/right cerebellar hemispheres and right/left cerebral hemispheres. For the CTC tractographies, a binarized mask of the superior cerebellar peduncle in MNI space from the Johns Hopkins University ICBM-DTI-81 white-matter labels atlas (Mori, S., Wakana, S., Van Zijl, P. C., & Nagae-Poetscher 2005; Wakana et al. 2007; Hua et al. 2018) was used as a waypoint, while the binarized contralateral cerebellar and cerebral hemispheres were set as exclusion masks (Example: CTC tractography between a left cerebellar ROI and a right cerebral ROI would entail a left superior cerebellar peduncle waypoint, a right cerebellar hemisphere exclusion mask, and a left cerebral hemisphere exclusion mask). For the CPC tractographies, a binarized mask of the middle cerebellar peduncle in MNI space from the same atlas was used as a waypoint, and the contralateral cerebellar and cerebral hemispheres were used as exclusion masks (Example: CPC tractography between a left cerebellar ROI and a right cerebral ROI would entail the middle cerebellar peduncle waypoint mask, a right cerebellar hemisphere exclusion mask, and a left cerebral hemisphere exclusion mask). We chose this method based on the prior tractography work of Palesi and colleagues (Palesi et al. 2015, 2017). The pons was not selected as an inclusion mask due to lack of a pons mask in standardized space through a standardized atlas. We also chose not to include the thalamus as a waypoint mask for the CTC pathway to keep the CTC and CPC tractography methods as similar as possible. Using probabilistic tractography, we recreated the same white matter pathways between motor (control) cerebellar ROIs and mentalizing cerebral ROIs. The number of streamlines for each path was obtained and used as a white matter integrity measurement.

## Results

### Functional domains in the Cerebellum

We first examined whether regions of the cerebellum have domain specific functions. At the group level, we extracted clusters of activation related to different task domains (Figure 1A) and local maxima (Figure 1B) for each task.

**Figure 1.**
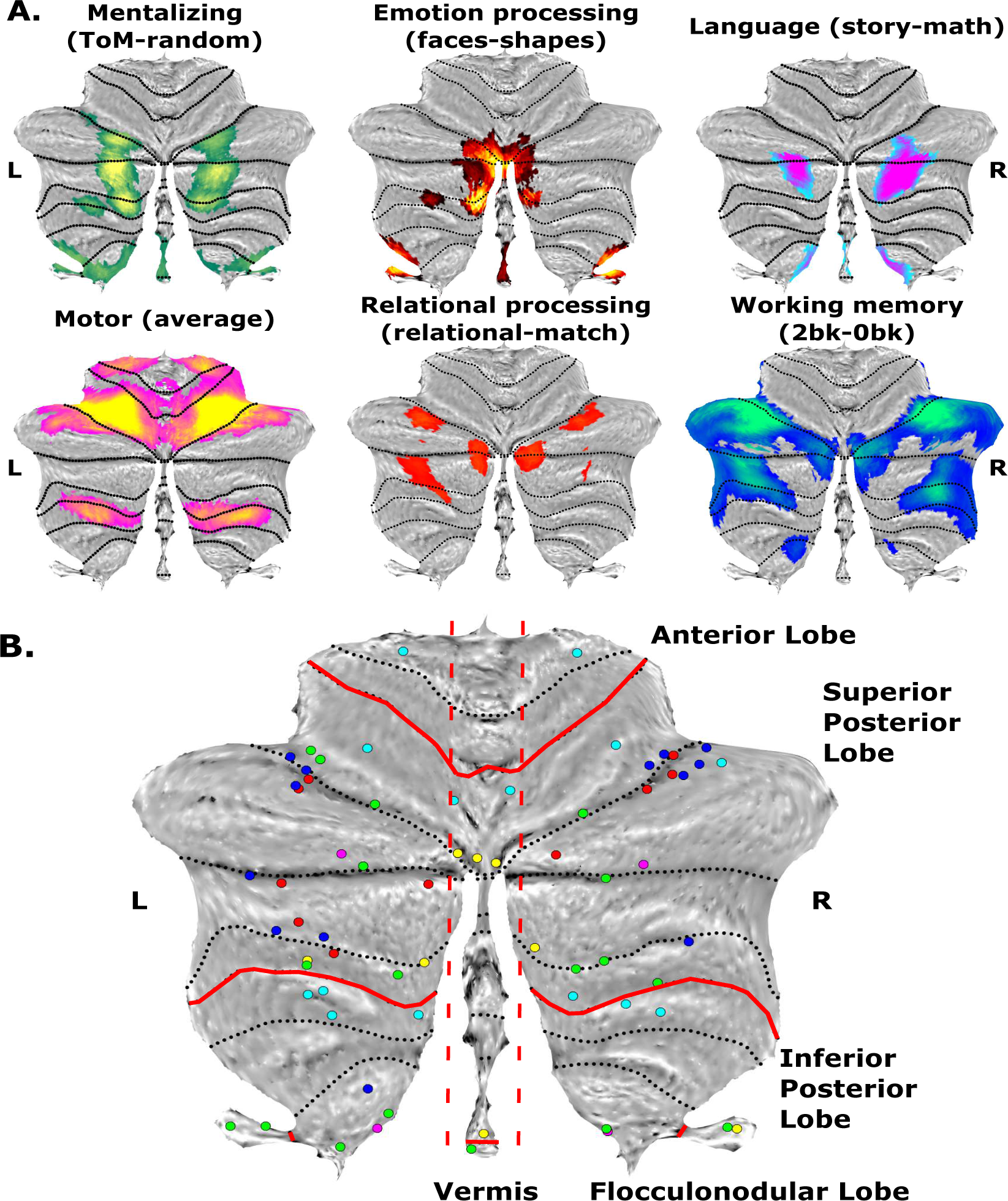
A. Group-level task clusters of activation for the six fMRI tasks. Activation colors indicate the particular task. L = left; R = right. B. Local maxima for each functional task. Each group of colored circles corresponds to a functional task. Yellow = emotion processing (L:5; R:4); Magenta = language processing (L:2; R:2); Turquoise = motor (L:7; R:7); Red = relational processing (L:6; R:4); Green = mentalizing (L:11; R:7); Blue = working memory (L:7; R:6). The total number of local maxima was 68 (L:38; R:30).

The Sørensen–Dice coefficient was used to calculate the overlap between the functional clusters of all tasks. Euclidean distances were then calculated to estimate the distances of local maxima within and between clusters. Lastly, at the individual level, we performed Wilcoxon signed-rank tests for each task pair to examine whether there was a statistical difference between them. All task contrasts generated activation clusters at the Cohen’s d > 0.5 threshold, except for the gambling task which was, subsequently, excluded from all analyses. Percentage of overlap between functional clusters from the tasks and cerebellar lobes are presented in Supplementary Table 1. Consistent with previous findings, the motor task was the only task that activated the anterior lobe (lobules I-VI), while emotion and mentalizing were the only tasks showing activation in the vermis (Gao et al. 2018; Albazron et al. 2019; Brady et al. 2019; Watson et al. 2019). Interestingly, mentalizing and emotion tasks also activated the flocculonodular lobe (lobule X), an area of the cerebellum known to regulate saccadic eye movements and balance (Cohen and Highstein 1972; Ito 1982; Schniepp et al. 2017). A detailed description of the overlap of each cluster of activation and each lobule is presented in Supplementary Table 2.

Percentage of overlap between task contrast activation clusters, calculated using the Sørensen–Dice coefficient, is presented in Table 1. The mentalizing and language clusters shared almost half of their voxels (45.17%), the mentalizing and emotion clusters share 18.66%, and mentalizing and working memory clusters shared 11.76%, while working memory shared voxels with motor and relational processing. The language and motor clusters were the only ones with zero overlap. The remainder task contrast cluster pairs had <10% overlap. To further examine the relationship between the task contrast activation clusters, we extracted the local maxima within each cluster and then calculated the mean Euclidean distance of the local maxima within each cluster and between clusters. 96.6% of Euclidean distances were > 8.5mm (whole sample of Euclidean distances *M* = 39.01mm, *SD* = 8.41mm) which means that only 78 pairs of local maxima were proximal. Out of the 78, only 58 were Euclidean distances of local maxima belonging to different clusters, bringing the percentage of local maxima that are proximal, yet belong to clusters of different task contrasts, to 2.55%. Despite shared activation, group-level Euclidean distances results indicate that overlapping clusters have distinct local maxima. These compelling findings at the group level require additional support from individual-level analyses to make a claim of generality vs specificity for cerebellar function.

**Table 1.** Percentage of overlap between task activation clusters. Percentages in bold are > 10%.

| Task | Emotion | Language | Motor | Relational<br>Processing | Mentalizing | Working<br>Memory |
| --- | --- | --- | --- | --- | --- | --- |
| Emotion |  |  |  |  |  |  |
| Language | 0.09 |  |  |  |  |  |
| Motor | 1.08 | 0 |  |  |  |  |
| Relational<br>Processing | 8.37 | 0.63 | 1.09 |  |  |  |
| Mentalizing | **18.66** | **45.17** | 5.50 | 2.89 |  |  |
| Working Memory | 7.50 | 0.41 | **20.99** | **19.61** | **11.76** |  |

At the single subject level, we extracted the averaged cerebellar activations for each task (i.e., across all cerebellar voxels with positive task contrast weights). Kolmogorov-Smirnov tests indicated that the task contrast-weights across subjects did not follow a normal distribution for all six tasks (*P’s* < .001; Supplementary Table 3) hence we used Spearman’s rho to examine the strength and direction of association between each task, and the Wilcoxon signed-rank test to examine whether there was a statistical difference in averaged cerebellar activation between each pair of tasks. We found significant but relatively weak positive correlations (*ρ’s* < .25) between emotion-language, emotion-relational, emotion-mentalizing, working memory-motor, working memory-language, and relational-working memory activations (Supplementary Table 4). Wilcoxon signed-rank tests showed that there was a statistically significant difference between contrast-weights of all task pairs (Supplementary Table 5) except for the relational-emotion pair. So despite the correlation of some task pairs, which can be attributed to the few voxels they share, the cerebellar activation of each task is significantly different from each other task. Together, the group-level and individual-level findings suggest that the cerebellum functions in a domain specific way with distinct cerebellar areas devoted to distinct cognitive functions.

### Functional Interactions between the Cerebellum and Cerebrum

Do mentalizing regions in the cerebellum functionally interact with mentalizing regions in the cerebrum? To measure this, we created 6mm spherical ROIs based on each mentalizing and motor cerebellar local maximum and on individual-subject peak activations of the mentalizing task for the ATL, DMPFC, PreC, TPJ, and VMPFC. We used PPI to examine the effective connectivity between the cerebellar and contralateral cerebral mentalizing areas and for the control analysis, cerebellar motor and contralateral cerebral mentalizing areas. This resulted in 110 mentalizing connections between the left cerebellum and right cerebrum (55 cerebellum-to-cerebrum e.g. L1 to R_ATL, L1 to R_DMPFC, 55 cerebrum-to-cerebellum e.g. R_ATL to L1, R_DMPFC to L1) and 70 connections between the right cerebellum and left cerebrum (35 CTC, 35 CPC). We extracted *Z*-scored *β*-weights for each pair of cerebellar to contralateral cerebral ROIs for each subject, averaged them, and at the group level we performed one-sample t-tests for each pair of ROIs to detect significant effective connections.

Each local maximum was represented in both cerebellar hemispheres and named accordingly (e.g. L1 – R1) with the exception of left hemisphere local maxima L7, L7a, and L8 which did not have the right cerebellar equivalent local maxima. In some cases, two local maxima were found in close proximity, in which case the extra local maximum was given the same name as the one near it, adding “a”. This resulted in the pairs L5-L5a (Euclidean distance = 8.485), L6-L6a (Euclidean distance = 7.21), L7-L7a (Euclidean distance = 4.90), and R1-R1a (Euclidean distance = 6.33). In order to understand whether these proximal ROIs are functionally distinct, we ran paired samples t-tests (or Wilcoxon signed-rank tests for connections that did not follow a normal distribution) on individual *β*-weights for all connections between the proximal ROIs (e.g. paired t-test comparing L5-ATL to L5a-ATL). Effective connectivity results for all R1 – R1a pairs was not significant but we found significant effective connectivity differences for 75% of proximal left ROI pairs (Supplementary Tables 6 & 7). These findings suggest that there is significantly different effective connectivity between mentalizing cerebellar and cerebral ROIs despite close proximity of the cerebellar local maxima.

Overall, we found more connections (Figures 2 and 3) and stronger effective connectivity between the left cerebellum and right cerebrum compared to vice versa. Local maxima on each of the left cerebellar lobes were connected to all five right mentalizing cerebral areas. On the left cerebellar hemisphere all local maxima ROIs were significantly connected to three or more right cerebral ROIs, with the exception of L8 on the flocculonodular node which was only connected to the right ATL and DMPFC. The right DMPFC, ATL, and TPJ displayed significant connections with most left cerebellar ROIs (10 out of 11 left cerebellar ROIs for the DMPFC and 9 out of 11 for the ATL and TPJ). The right cerebellar hemisphere displayed a different pattern of connectivity. With the exception of R1 and R1a, which combined were significantly connected to all left cerebral ROIs, all other right cerebellar ROIs were significantly functionally connected to either one or two left cerebral ROIs. The left TPJ was the brain area which displayed significant effective connectivity with all right cerebellar ROIs, except R4. Combining the significant connections of all the right cerebellar ROIs, we find that the right superior posterior, inferior posterior, and flocculonodular lobes display significant connectivity to all five left mentalizing brain areas. The right superior posterior cerebellar lobe is significantly connected to all five left cerebral brain areas when we combine the significant connectivity results of all its cerebellar ROIs (R1-R4), but the right inferior posterior lobe is only connected to the left ATL and TPJ, and the right flocculonodular lobe is only connected to the left TPJ (Figure 3).

**Figure 2.**
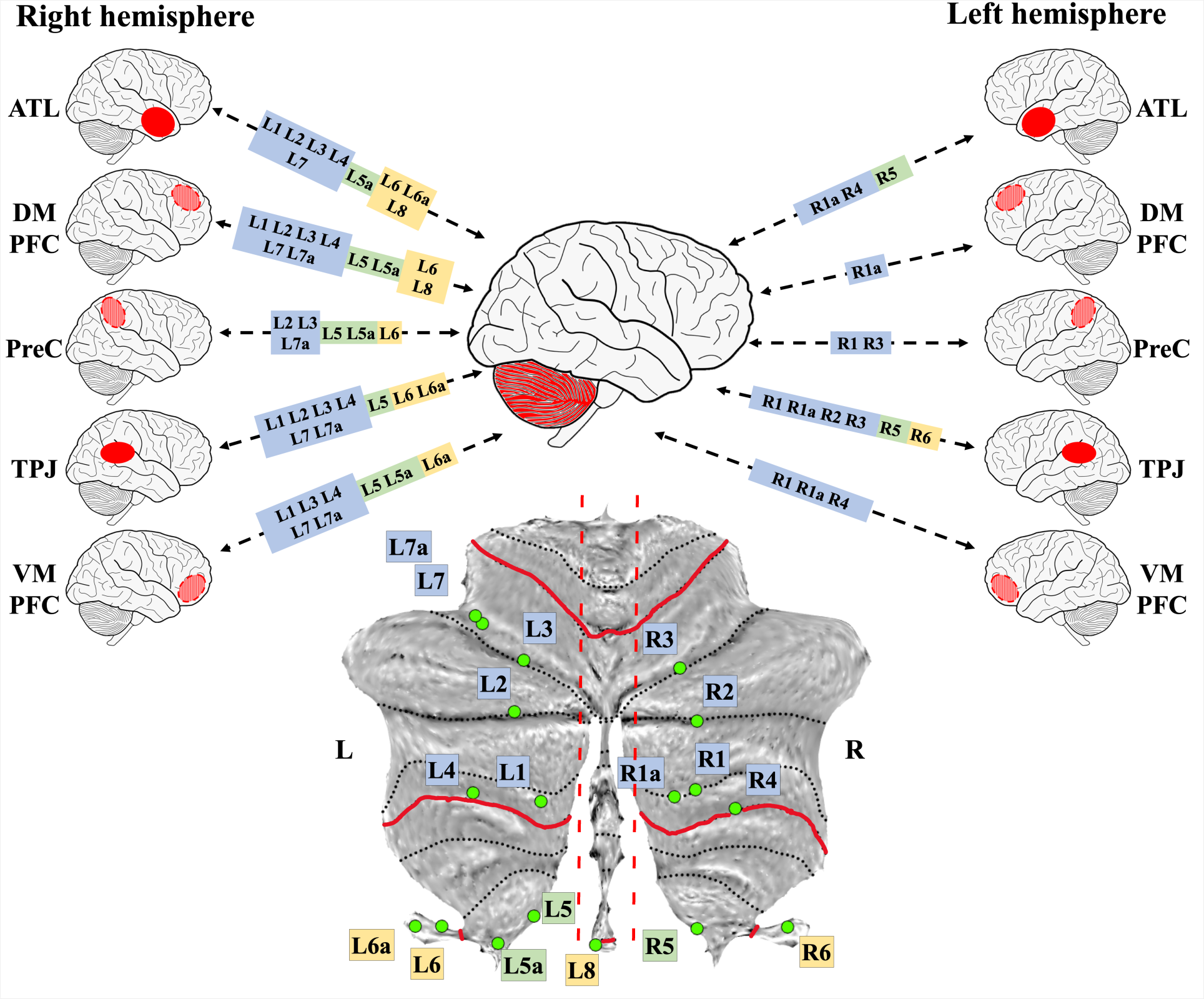
Effective connectivity between cerebellar and cerebral mentalizing ROIs. The color of each square denotes its cerebellar lobe (e.g. blue = superior posterior lobe; green = inferior posterior lobe; yellow = flocculonodular lobe). If a local maximum was found in both cerebellar hemispheres, it was named accordingly (e.g. L1 – R1). If two local maxima were found in close proximity, the extra local maximum was denoted by the letter “a”. L=left; R=right. ATL = anterior temporal lobe; DMPFC = dorsomedial prefrontal cortex; PreC = precuneus; TPJ = temporoparietal junction; VMPFC = ventromedial prefrontal cortex.

**Figure 3.**
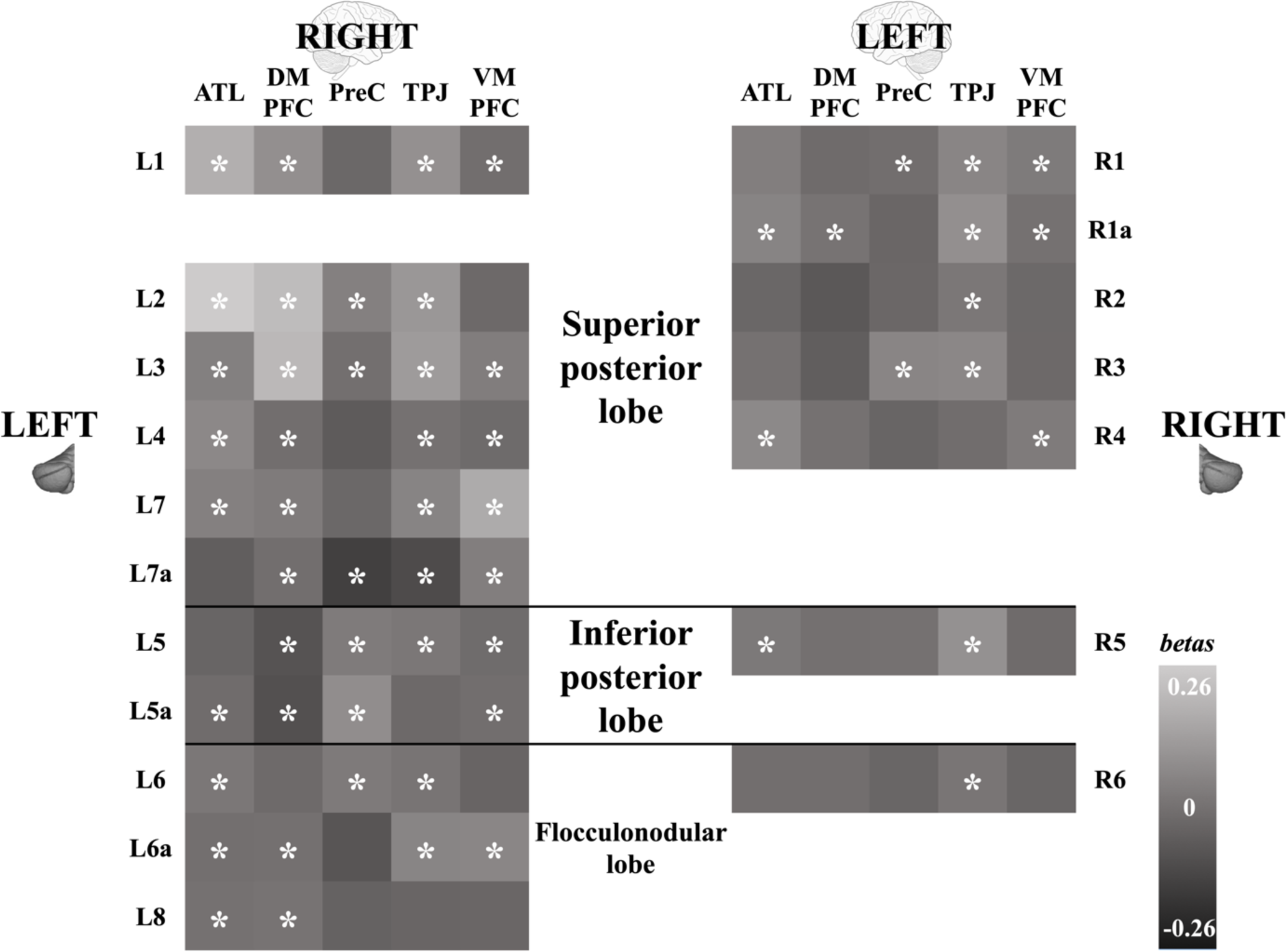
Effective connectivity significance between cerebellar and cerebral mentalizing ROIs. Stars indicate significant connections. ATL = anterior temporal lobe; DMPFC = dorsomedial prefrontal cortex; PreC = precuneus; TPJ = temporoparietal junction; VMPFC = ventromedial prefrontal cortex.

To test the specificity of these functional connections, we ran PPI analyses between cerebellar motor ROIs and the contralateral mentalizing cerebral ones and performed the same analyses as above. We found that, with minor exceptions, cerebellar motor to cerebral mentalizing effective connections were not significant (*P’s* > .06; Supplementary Table 8). Additionally, we ran a two-sample binomial proportion test between the mentalizing and motor effective connections for both cerebellar hemispheres and found that the proportion of mentalizing-mentalizing significant connections differed significantly from the proportion of motor-mentalizing connections with significantly more mentalizing-mentalizing connections in both cerebellar hemispheres (Left cerebellum to right cerebrum: *z* = 4.2, *P* < .0001, Right cerebellum to left cerebrum: *z* = 2.7, *P* < .01).

These results indicate that there is a clear functional connection between regions sensitive to mentalizing in the cerebellum and regions sensitive to mentalizing in the cerebrum. There is some lateralization of function with the left cerebellar lobes communicating with all right mentalizing cerebral areas while the right cerebellum displays more limited connectivity. In addition, all left cerebellar local maxima were functionally connected to the right DMPFC, but only one right local maximum in the cerebellum was (weakly) connected to the left DMPFC. Notably, local maxima in both cerebellar hemispheres showed strongest connectivity to the TPJ.

### Structural Connections between the Cerebellum and Cerebrum

Are cerebellar mentalizing areas structurally connected to cerebral mentalizing areas? To test this, we ran probabilistic tractography to reconstruct the CTC and CPC white matter pathways. We used the same 6mm spherical ROIs from the PPI analyses as seeds and targets for the tractographies, the superior cerebellar peduncle as a waypoint for the CTC white matter pathway, and the middle cerebellar peduncle for the CPC white matter pathway (Palesi et al. 2015, 2017). We then extracted streamline counts for each pathway (Supplementary Tables 9 and 10).

Overall, the right/left CTC and right/left CPC pathways showed similar connectivity patterns and average streamline counts (Figure 4). We combined the individual average streamline counts to create group average streamline counts for the superior posterior, inferior posterior, and flocculonodular lobes for the right and left CTC and CPC pathways (Table 2). We used Spearman’s rho (the white matter streamline counts did not follow a normal distribution; *P’s* < .001; Supplementary Table 11) to examine the strength and direction of association between left and right CTCs and CPCs, and the Wilcoxon signed-rank test to examine whether there was a statistical difference between them. For both the CTC and CPC pathways we found that the right superior posterior streamline count is correlated to the left superior posterior, inferior, and flocculonodular lobes but it shows greater correlation to the left superior posterior lobe. The same applies for the left superior posterior streamline counts and the inferior posterior and flocculonodular lobe streamline counts for both the right and left cerebellar hemispheres (Supplementary Tables 12 & 13). Wilcoxon signed-rank tests showed that there was a statistically significant difference in streamline counts between each lobe pair (e.g. left superior posterior – right superior posterior) for CTC and CPC tracks, with the majority of subjects having significantly larger CTC streamline counts in the right superior posterior lobe rather than the left, but significantly smaller streamline counts in the right inferior and flocculonodular lobes compared to the respective left. The exact same pattern was observed in the CPC tracks. We also found that the overall left CTC streamline count was significantly larger compared to the right CTC streamline count and the same was true for the CPC streamline counts. Additionally, we found that CTC streamline counts (from the left and right cerebellar hemispheres) were significantly larger compared to the ipsilateral CPC streamline counts (Supplementary Table 14). CTC pathways from both the left and right cerebellar hemisphere and towards all five contralateral mentalizing cerebral ROIs displayed the exact same streamline count pattern (DMPFC>VMPFC>PreC>ATL>TPJ). The CPC pathways displayed a different pattern. In the CPC pathways projecting to the left cerebellar hemisphere, the right TPJ was the brain area with the second largest streamline count towards all left cerebellar ROIs, except for L8. Also, in the CPC pathways projecting to the right cerebellar hemisphere, the left VMPFC displayed disproportionally smaller streamline counts towards all right cerebellar ROIs compared both to the left CPC pattern and both CTC patterns (Figure 4; Supplementary Tables 9 & 10). Overall, the largest streamline count was both to and from the DMPFC.

**Figure 4.**
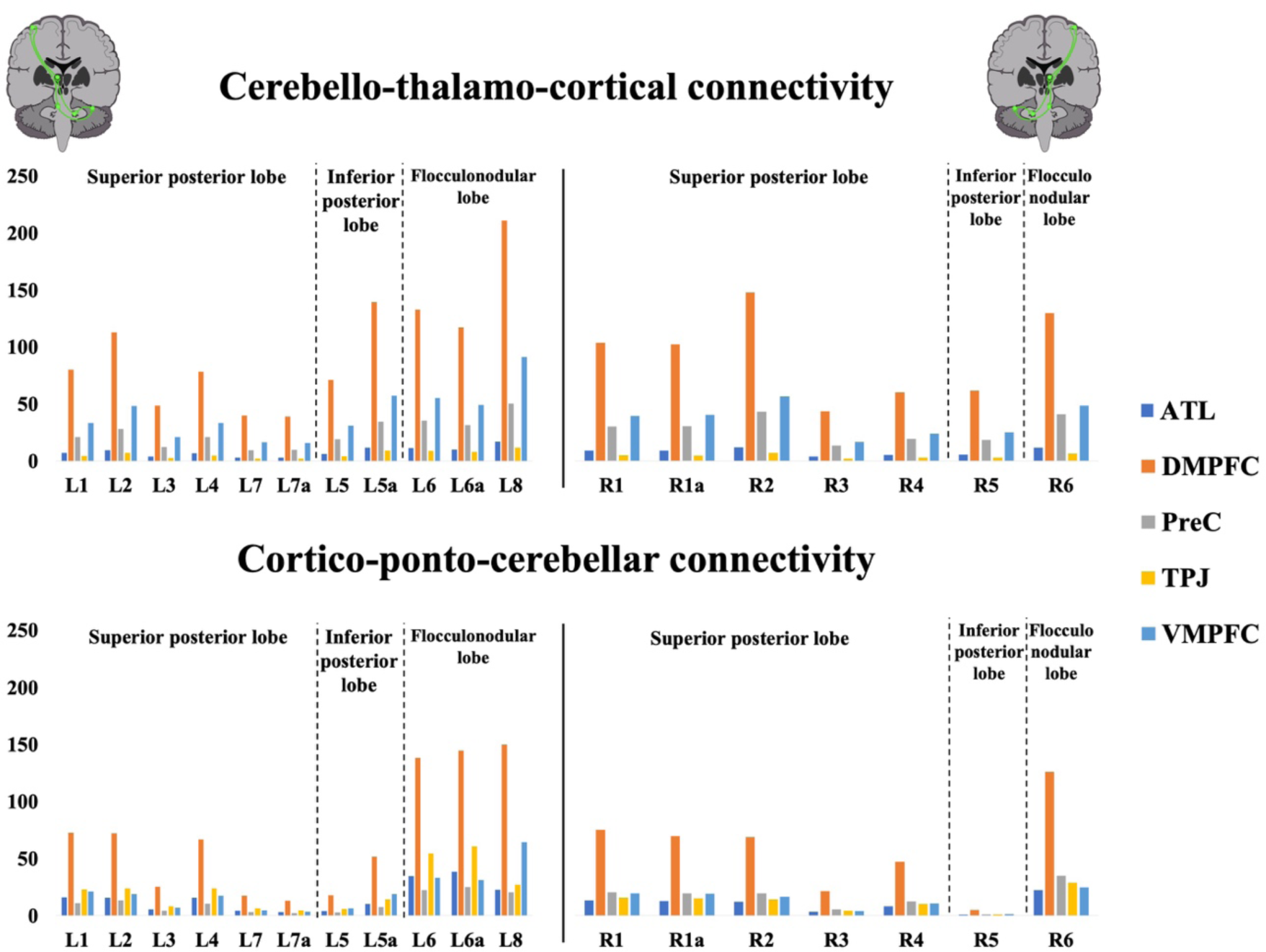
Mean streamline counts for each cerebello-thalamo-cortical and cortico-ponto-cerebellar white matter tract. The left column contains streamline counts seeded from mentalizing ROIs from the left cerebellum. The right column contains streamline counts seeded mentalizing ROIs from the right cerebellum. ATL = anterior temporal lobe; DMPFC = dorsomedial prefrontal cortex; PreC = precuneus; TPJ = temporoparietal junction; VMPFC = ventromedial prefrontal cortex.

**Table 2.** Average cerebello-thalamo-cortical (CTC) and cortico-ponto-cerebellar (CPC) streamline counts per cerebellar hemisphere and lobe. The average streamline counts of whole mentalizing CTC and CPC pathways are in bold.

|  | Streamline count |  |  | Streamline count |  |
| --- | --- | --- | --- | --- | --- |
|  | Mean | SD |  | Mean | SD |
| **Left CTC** | **35.52** | **(23.07)** | **Right CTC** | **32.76** | **(22.77)** |
| Left superior posterior lobe | 24.27 | (18.79) | Right superior posterior lobe | 32.26 | (24.54) |
| Left inferior posterior lobe | 38.42 | (30.75) | Right inferior posterior lobe | 22.04 | (23.65) |
| Left flocculonodular lobe | 56.10 | (34.66) | Right flocculonodular lobe | 45.99 | (32.48) |
| **Left CPC** | **28.04** | **(20.64)** | **Right CPC** | **21.74** | **(17.01)** |
| Left superior posterior lobe | 17.82 | (13.45) | Right superior posterior lobe | 20.89 | (16.80) |
| Left inferior posterior lobe | 13.96 | (12.94) | Right inferior posterior lobe | 1.86 | (2.71) |
| Left flocculonodular lobe | 57.84 | (44.96) | Right flocculonodular lobe | 45.87 | (48.06) |

To examine the specificity of these white matter connections we ran probabilistic tractographies between the motor cerebellar areas and contralateral mentalizing cerebral ones. Results showed that the white matter pathways from the motor cerebellar areas to the mentalizing cerebral areas had low streamline counts (Supplementary Table 15) and Wilcoxon signed-rank test comparing the mentalizing-mentalizing and motor-mentalizing connections showed that the mentalizing-mentalizing white matter pathways had significantly larger streamline counts in each cerebellar lobe but also overall for the CTC and CPC (*P’s* < .001; Supplementary Table 16).

These results indicate that there are structural connections between the mentalizing cerebellar and cerebral areas. The feedforward (CTC) and feedback (CPC) pathway streamline counts between the right and left cerebellar lobes differed significantly. The right cerebellar superior posterior lobe had a larger streamline count than its left counterpart, but the right inferior and flocculonodular lobes had smaller streamline counts compared to their left counterparts. Additionally, there was a left lateralization in white matter connectivity with the overall left CTC streamline count being significantly larger compared to the right CTC streamline count, and the same being true for the CPC streamline counts. Of all ROIs in the classic mentalizing network, the DMPFC had the largest streamline count in both directions.

## Discussion

In this study we explored whether the cerebellum plays a role in mentalizing. We used a large-scale dataset and mapped out the cerebellum’s functional organization, functional connectivity, and structural connectivity. First, we investigated whether there is functional specificity in the cerebellum for mentalizing. To assess this, we looked at activation overlap between mentalizing activations and five other task activations: emotion processing, language, motor, visual relational processing, and working memory. There was sizeable overlap between mentalizing, emotion, and language. The mentalizing-emotion overlap occurred in the vermis and bilateral flocculonodular lobes (Figure 1) and was anticipated. In non-human animals the vermis and flocculonodular lobes are structurally connected to the amygdala, hippocampus, septum, and hypothalamus (Heath and Harper 1974; Snider et al. 1976; Hu et al. 2008) thus these regions have long been thought of as the limbic system of the cerebellum (Schmahmann, 1994). Vermal stimulation in mice, cats, and primates leads to fire pattern modulation in these structures (Babb, Mitchell, & Crandall, 1974; Berman, 1997; Bobée, Mariette, Tremblay-Leveau, & Caston, 2000; Zanchetti & Zoccolini, 1954) and aggression amelioration in human patients (Heath et al. 1979). Damage to the vermis can lead to atypical social and emotional behaviors (Schmahmann 1991; Pollack 1997; Schmahmann and Sherman 1998; Levisohn et al. 2000; Tavano et al. 2007; Manto and Mariën 2015) and neuroimaging studies have confirmed that the vermis and flocculonodular lobes are active during emotion learning or change of affect tasks (Lane et al. 1997; Beauregard et al. 1998; Gündel et al. 2003) and social processing (Van Overwalle et al. 2014; Guell, Gabrieli, et al. 2018). This overlap also makes sense given that these tasks contained overlapping features and processes. For instance, the emotion task involved some degree of implicit mentalizing because participants were required to attribute emotion to faces (Wang et al. 2020). One could also argue that most language tasks that contain any narrative require mentalizing. The task in the HCP dataset actually has an unusually high mentalizing load since the stimuli were variants of Aesop’s fables, which involved animals or humans interacting in social situations (Binder et al. 2011). Whether language tasks that are less social – for instance, single word processing – would show overlap with mentalizing activations in the cerebellum is not known. Task activation clusters and their overlap are heavily dependent on thresholding decisions. Guell et al. (2018) also used the HCP S900 dataset to examine the task representations on the cerebellum but they analyzed the 2mm smoothed fMRI data (here we used 4mm smoothed data). Although their task clustering results are similar to ours, the cluster overlaps were smaller, with overlap found solely between the mentalizing and language tasks. They conclude that the cerebellum has domain-specific representations of different kinds of cognition and emotion. We believe this is a reasonable conclusion, but we wanted to provide additional evidence which would strengthen the claim to a domain-specific cerebellar role. The task cluster overlaps led us to explore in more depth the relationship between the activation clusters. Our Euclidean distance analysis showed that each cluster had distinct local maxima. We also analyzed the strength of cerebellar activation for each task at the individual level and found that each task elicited different degrees of cerebellar activations across subjects. Overall, our results provide support for the hypothesis that the cerebellum has domain-specific functions.

Second, we reasoned that if mentalizing clusters in the cerebellum truly play a role in mentalizing behavior, they should show effective connectivity with known mentalizing regions in the cerebrum. We tested this and found that (1) there was strong effective connectivity with all mentalizing regions in the cerebral cortex (Figure 2); and (2) there are laterality differences, with clusters in the left cerebellar hemisphere displaying more and stronger effective connections to the right cerebral mentalizing areas, compared to the right cerebellar – left cerebral hemisphere. These results are consistent with the right hemisphere bias of the mentalizing network in the cerebrum (Wang et al. 2020), with multiple studies showing both task activity and within-cerebrum connectivity lateralized in the right ATL (Gainotti 2007; Olson et al. 2007, 2013; Acres et al. 2009; Rice et al. 2015) and right TPJ (Saxe and Wexler 2005; Van Overwalle and Baetens 2009; Saxe 2010; Santiesteban et al. 2015; Karolis et al. 2019). In our study, the ATL displayed this lateralization with the right ATL being significantly connected to all but two left mentalizing cerebellar ROIs. Interestingly, the TPJ did not display this lateralization. The role of the TPJs in mentalizing is debated with several studies showing unilateral right TPJ activation in mentalizing (see above), but others showing bilateral involvement (Gallagher et al. 2000; Van Overwalle 2009; Jenkins and Mitchell 2010; Bzdok et al. 2012; Schurz et al. 2014; Molenberghs et al. 2016). Despite this debate, we attribute the bilateral TPJ activation to the fact that the mentalizing task involved videos of socially meaningful biological motion. In their study on the underlying neural responses of anthropomorphism in autism spectrum disorder and typically developing individuals, Ammons et al. (2018) found bilateral TPJ activation during observation of social movement regardless of the type of agent (human figure or geometrical shape) in both autistic and typically developing individuals. Furthermore, in our study we used a broader definition of the TPJ which could include parts of the IPL, known to be involved in perspective taking, and portions of the pSTS, involved in biological motion perception. We believe that the nature of the stimuli used to activate the mentalizing areas, combined with our selection method of the TPJ ROI, could explain the significant bilateral cerebellar effective connectivity of almost all cerebellar local maxima and the contralateral TPJs.

Third, we examined the structural connectivity profile using probabilistic tractography between cerebellar and cerebral mentalizing areas. The cerebellum is connected to the cerebral cortex through complex ouroboros loops. The CTC pathway sends projections from the cerebellum to the cortex, then that cortical region projects back to the same cerebellar region through the CPC pathway. Anatomical tracer studies in non-human animals have shown that cerebello-cerebral structural connectivity is different between primates and other vertebrates, with primate cerebellar output fibers reaching the frontal association cortex (Ito 1984; Strick et al. 2009; Stoodley and Schmahmann 2010). Only a few studies have employed diffusion-weighted imaging techniques and tractography to confirm in-vivo the existence of white matter connections in humans between the cerebellum and the cerebral cortex and explore their quality. Jissendi et al. (2008) were the first to isolate the complete cerebellar projections to prefrontal and posterior parietal cortices using tractography, albeit in a small sample and using rudimentary techniques. Similarly, Sokolov et al. (2014) and Keser et al. (2015) delineated these tracts in small samples. Karavasilis et al. (2019) confirmed the results by Keser et al. (2015) by replicating their study with a larger number of participants *(N* = 60). All of the above findings are promising evidence that the cerebellum is anatomically connected with non-motor cerebral areas, but the in vivo anatomical evidence for specific cerebellar and cerebral mentalizing areas is still extremely limited.

We recreated the CTC and CPC white matter pathways using probabilistic tractography. There were three primary findings. First, we found that the CTC pathways had greater streamline counts compared to the CPC pathways. Second, there was some lateralization such that there were relatively larger streamline counts in the left cerebellar CTC and CPC pathways compared to the right. This result matches nicely with the lateralization observed in our effective connectivity results. Third, white matter streamline counts were dramatically higher between mentalizing cerebellar regions and the DMPFC, followed by the VMPFC (Figure 4). This is only partially consistent with the effective connectivity findings since the TPJ was the cerebral area with the most, significant effective connections in our analysis, followed by the DMPFC, the ATL, and then the VMPFC. There is a well-known quantitative mismatch between structural and functional connectivity (Huang and Ding 2016; Suárez et al. 2020). We note that there is solid “ground truth” for the DMPFC and VMPFC findings since histology methods have revealed fiber connections between the cerebellum and non-motor regions of the frontal lobe (Ito 1984; Leiner et al. 1993; Middleton and Strick 1997, 2001; Schmahmann and Pandya 1997; Dum and Strick 2003; Kelly and Strick 2003; Clower et al. 2005); whether they exist in the TPJ, ATL and precuneus is not known. As such we feel that the structural connectivity and functional connectivity results between the cerebellum and the DMPFC and VMPFC stand on firm ground. In contrast, we feel that more work is needed to understand whether structural pathways exist between the cerebellum and parietal and temporal cortices. Even if direct pathways via the CTC/CPC are not present, other polysynaptic routes exist. For instance, the uncinate fasciculus connects the ATL to the VMPFC (Von Der Heide et al. 2013) and from there, the CTC pathway begins.

Our findings add to a small but growing literature linking the cerebellum to social cognition. However, a key unanswered question is “what is the cerebellum’s mechanistic or computational process?”. In the motor literature there is agreement that the cerebellum contributes to the regulation of the rhythm, rate, accuracy, and force of movements (for a review see Manto et al. 2012). Similarly, it is considered that the cerebellum regulates the capacity, speed, appropriateness and consistency of mental processes. The theory that the cerebellum maintains behavior around a homeostatic baseline is termed the “dysmetria of thought theory” (Schmahmann 1998) and it is analogous to the dysmetria of movement theory. The latter is presented with overshooting and lack of control in the motor system. In the dysmetria of thought theory, these are equated with unpredictability in social interactions, a mismatch between reality and perceived reality, and unsuccessful, illogical efforts to correct errors in behavior and thought (Schmahmann 2019). It is clear that dysmetria of movement is matched with dysmetria of thought and this match in functions is explained by the theory of the Universal Cerebellar Transform (Schmahmann, 2000, 2001, 2004) according to which there is a unique cerebellar computation which is applied to all its functions (movement- and cognition-related) (Schmahmann 2000, 2001, 2004; Guell, Gabrieli, et al. 2018) due to its uniform cortical cytoarchitectonic organization (Ito, 1993; Voogd & Glickstein, 1998). However, it is also possible that the cerebellar circuitry performs multiple diverse computations, given that the cerebellum appears with striking functional heterogeneity (Diedrichsen et al. 2019). The most prominent theory about the nature of the cerebellar computation posits that the cerebellum is encoding internal models that reproduce the dynamic properties of body parts for controlling these body parts without any sensory feedback, and for mental processes, that it encodes internal models which reproduce the essential properties of mental representations in the cerebral cortex (Ito 2008). In a recent paper, Van Overwalle et al (2019) formed a hypothesis positing that, when it comes to social cognition, the cerebellum acts as a “forward controller” predicting how actions by the self and other people will be executed, what are the most likely responses to these actions, and what is the typical sequence of these actions. So in the realm of motor behavior, the cerebellum calibrates the kinematics of movement, potentially by comparing an internal model to reality, computing an error, and making online adjustments. It has been proposed that something similar occurs for social cognition but direct tests of this are rare.

Does cerebellar damage cause social cognition or mentalizing deficits? We recently reviewed the lesion literature (Olson et al. 2021) and found that in infants the answer is yes, in children the answer is “sometimes”, and in adults the answer is “rarely.” Whether the larger and more consistent effects observed in children are due to their developing brain, the relatively larger scale of the cerebellar damage, or the way in which social cognition is measured, is not known. However two features of the adult studies deserve inspection. First, the tasks used in adult lesion studies may be ill-suited for the computational processes that occur in the cerebellum. Many mentalizing tasks are performed in a nearly error-free manner by neurologically normal adults. A Heider and Simmel-type task, such as the one used here, may be one of the best tasks for evoking cerebellar activations because the stimuli are novel and the shapes move in unpredictable ways, which triggers a constant stream of rapid predictions and error signals. At the same time, there is a social narrative that is being played out, so internal models are activated. However, most lesion studies use Reading the Eyes in the Mind (Baron-Cohen et al. 2001; Fernández-Abascal et al. 2013) or some variant of false-belief stories. Second, the fact that adults with any type of cerebellar damage are included in these studies adds noise to the measure. As we saw in the present study, mentalizing clusters are limited to a small region of the posterior cerebellum so individuals with lesions that preserve this area should not be expected to exhibit social deficits or problems performing mentalizing tasks.

### Limitations and Future Directions

First, connectivity analyses are inherently vulnerable to conceptual and methodological issues (Buckner et al. 2013) and their results depend on various factors (e.g. ROI selection methods, stimuli features, thresholding choices) (Uddin et al. 2008; Turchi et al. 2018). Accordingly, our findings should be interpreted with caution. Furthermore, diffusion MRI tractography has received criticism for being a method with high rates of false positives (Thomas et al. 2014; Reveley et al. 2015; Maier-Hein et al. 2017). Regardless, functional and diffusion MRI have provided valuable insight into the anatomical and functional organization of the mentalizing network, and as both techniques improve, we hope other researchers replicate and extend our cerebellar findings.

Second, the task set included in the HCP dataset is limited. Whether other tasks with mentalizing demands – sarcasm processing, deception, or false beliefs – engage the same regions of the cerebellum and are functionally and structurally connected with the greater mentalizing network in the cerebrum needs to be explored by future researchers. Related to this question, it is unclear what aspect of mentalizing is driving this network. There are different types of mentalizing: conceptual (e.g. false-belief), perceptual (e.g. face-based), affective (e.g. empathy, moral reasoning), cognitive (e.g. perspective-taking), and motion/motoric (e.g. Heider-Simmel biological motion trajectory understanding) (Luyten et al. 2020). Is the cerebellum implementing the same computational mechanism regardless of the particular mentalizing task? This seems implausible, given the wide variety of computational mechanisms used to perform the various tasks that fall into this category. It has recently been proposed that mentalizing tasks with a sequential structure – like the one used in this study – are the only ones linked to cerebellar function (Van Overwalle et al. 2019), potentially because they draw more heavily on predictive modeling computations (Gonzalez and Chang 2019) linked to cerebellar function more generally. These hypotheses need to be further explored in the future.

## Conclusions

This multimodal neuroimaging study investigated the structural and functional connectivity between mentalizing cerebellar and cerebral areas. We found that the cerebellum functions in a domain specific way with distinct cerebellar areas devoted to distinct cognitive functions, among which is mentalizing. Furthermore, we mapped out the effective connectivity between the mentalizing cerebellum and cerebrum and found cerebellar hemispherical differences in cerebello-cerebral effective connectivity. The left cerebellar hemisphere displayed more and stronger significant effective connections to the right cerebral mentalizing areas. Additionally, we recreated the CTC and CPC pathways using probabilistic tractography and found that the CTC and CPC pathways from and to the left cerebellum had larger streamline counts compared to the right CTC and CPC pathways. Lastly, we found that the bilateral CTC pathways had greater streamline counts compared to the CPC pathways. All the above findings suggest that regions of the posterior cerebellum play a key role in mentalizing abilities.

## Acknowledgments

We would like to thank Geoffrey Wright, Vishnu Murty, Jason Chein, Huiling Peng, and Nico Dosenbach for theoretical input at the dissertation phase that benefited this publication, and Dr. Axel Kohlmeyer for assistance with high performance cluster computing.

