## Supplementary Material for "The social cerebellum: A large-scale investigation of functional and structural specificity and connectivity"

Human Connectome Project protocol

The HCP protocol includes acquisition of structural MRI (0.7mm isotropic voxels), resting-state fMRI (rfMRI) (2mm isotropic voxels, four runs of 1200 volumes, TR = 720mm, 14m33sec per run), tfMRI (2mm isotropic voxels, TR = 720ms, tasks: working memory, gambling, motor, language processing, relational processing, social cognition, emotion processing), and dMRI (1.25mm isotropic voxels, three shells of b = 1000, 2000, and 3000s/mm^2,^, 90 diffusion-weighting directions acquired with right-to-left and left-to-right phase encoding) in a customized Siemens 3T “Connectome Skyra” scanner. Participants also received extensive behavioral testing (Barch et al. 2013) Additional measures that were obtained included, among others, demographics information, health measures, sleep pattern, drug screening.

In-scanner Tasks

Participants were assessed in seven major domains: 1) social cognition (mentalizing); 2) motor (visual, motion, somatosensory, and motor systems); 3) gambling; 4) working memory/cognitive control systems and category specific representations; 5) language processing (semantic and phonological processing); 6) relational processing; and 7) emotion processing (Barch et al. 2013). Detailed information on the tasks used to assess each domain can be found in the respective articles: The motor task was an adaptation of the task developed by Buckner and colleagues (Buckner et al. 2011; Yeo et al. 2011) and included finger, toe, and tongue movements; the gambling task was a modified version of the task developed by Delgado et al. (2000) and included guessing whether a symbol represented a number bigger or smaller than 5 and receiving feedback on the guess; the working memory task was a 2-back task with blocks of trials consisting of images of faces, places, tools, and body parts; the language processing task (henceforth “language”) was developed by Binder et al. (2011) consisted of brief auditory adapted stories from Aesop’s fables followed by a two-alternative forced-choice question asking participants about the topic of the story; the relational processing task (henceforth “relational”) was an adaptation of a task developed by Smith, Keramatian, & Christoff (2007) in which participants were asked to match a top item to a bottom one based on two different dimensions; and the emotion processing task (henceforth “emotion”), adapted from the one developed by Hariri et al. (2006), in which participants were asked to match the emotion of two faces presented at the bottom of the screen to that of a face presented at the top. Lastly, in the main task of interest, the social cognition task (henceforth “mentalizing”), participants viewed 20 s video clips of geometrical shapes that either interacted with each other or moved purposelessly on the screen. The videos were developed by either Castelli et al. (2000) or Wheatley et al. (2007) and have been validated as a measure of mentalizing given evidence that they generate task related activation in brain regions associated with mentalizing with reliable results across subjects (Castelli et al. 2000, 2002; Wheatley et al. 2007; White et al. 2011; Barch et al. 2013). Both the emotion and the mentalizing task require some degree of mentalizing. The mentalizing task asks participants to implement mental state attributions when watching the geometrical shapes interact, hence the mentalizing is intentional. On the other hand, the emotion task has a far smaller degree of mentalizing as participants only implicitly attribute mental states to the faces they view (Van Overwalle and Vandekerckhove 2013; Kliemann and Adolphs 2018).

Image Preprocessing

The imaging data used in this article were the “minimally preprocessed” included in the WU-Minn HCP Consortium S900 Release (WU-Minn HCP Consortium 2015). The dMRI data preprocessing included echo planar imaging (EPI) distortion, eddy-current-induced distortion, and subject motion correction, gradient nonlinearity correction, normalization of the b0 image intensity across runs, and registering the mean b0 volume to a native T1 volume. The tfMRI data had undergone spatial artifact/distortion correction, cross-modal registration, and spatial normalization to MNI space. Additionally, we further processed the dMRI data with FSL’s BEDPOSTX multi-shell, ball and zeppelins model (Hernández et al. 2013; Sotiropoulos et al. 2016) to model white matter fiber orientations and crossing fibers, and removed motion artifacts from the tfMRI data using ICA-AROMA (Griffanti et al., 2014; Pruim, Mennes, Buitelaar, & Beckmann, 2015; Pruim et al., 2015). All fMRI data were spatially smoothed at 4mm.
