## Supplementary Tables for "The social cerebellum: A large-scale investigation of functional and structural specificity and connectivity"

**Supplementary Table 1.** Task contrast activation clusters and cerebellar lobe percent overlap. The percentages in each row do not add up to 100% as part of the activations were categorized as “Unknown”. Percentages in bold are > 10%.

| Task | Cerebellar  lobe | Anterior | Posterior | Flocculonodular | Vermis |
| --- | --- | --- | --- | --- | --- |
| Emotion | | 0.002 | **62.54** | 9.07 | **18.56** |
| Language | |  | **92.83** | 0.01 | 0.38 |
| Motor | | **17.04** | **71.35** | 0.004 | 1.96 |
| Relational | |  | **95.98** |  | 0.13 |
| Mentalizing | | 0.0008 | **87.96** | 1.91 | 2.98 |
| Working Memory | | 0.03 | **91.14** | 0.02 | 0.73 |

**Supplementary Table 2.** Task contrast activation clusters and cerebellar lobule percent overlap. Percentages > 10% are in bold.

| Cerebellar lobes | Cerebellar lobules | Emotion | Language | Motor | Relational  Memory | Mentalizing | Working  Memory |
| --- | --- | --- | --- | --- | --- | --- | --- |
| Left anterior lobe | Left I-IV | > 0.01 |  | 1.60 |  |  | > 0.01 |
|  | Left V |  |  | 5.34 |  | > 0.01 | 0.02 |
| Left superior  posterior lobe | Left VI | 9.95 |  | **19.90** | 5.74 | 5.14 | 8.88 |
|  | Left Crus I | 5.14 | 7.70 | 4.95 | **21.97** | **13.82** | **22.68** |
|  | Left Crus II | **25.41** | 9.61 | 0.09 | **30.41** | **20.40** | **12.61** |
|  | Left VIIb | 7.85 | 0.01 | 2.95 | 7.00 | 7.68 | 5.33 |
| Left inferior  posterior lobe | Left VIIIa | 0.19 |  | 4.08 | > 0.01 | 1.10 | 0.44 |
|  | Left VIIIb | 0.30 | 0.01 | 0.02 |  | 0.11 | 0.18 |
|  | Left IX | 0.39 | 1.41 |  |  | 4.34 | 0.75 |
| Left  flocculonodular lobe | Left X | 4.77 |  |  |  | 1.05 | 0.01 |
| Right  anterior lobe | Right I-IV |  |  | 3.32 |  |  |  |
|  | Right V | > 0.01 |  | 6.79 |  | > 0.01 | 0.01 |
| Right superior  posterior lobe | Right VI | 3.09 | 1.13 | **20.93** | 4.66 | 3.86 | 7.71 |
|  | Right Crus I | 1.58 | **29.03** | 6.18 | **24.71** | **11.04** | **17.61** |
|  | Right Crus II | 6.68 | **40.12** | 0.33 | 1.49 | **16.49** | 7.33 |
|  | Right VIIb | 1.40 | 0.19 | 4.27 |  | 2.97 | 6.36 |
| Right inferior  posterior lobe | Right VIIIa | 0.09 |  | 7.26 |  | 0.04 | 1.25 |
|  | Right VIIIb | 0.27 | 0.01 | 0.38 |  | 0.06 | 0.01 |
|  | Right IX | 0.20 | 3.63 | 0.01 |  | 0.91 |  |
| Right  flocculonodular lobe | Right X | 4.30 | 0.01 | > 0.01 |  | 0.86 | 0.01 |
| Vermis | Vermis VI | **10.86** |  | 1.93 | 0.03 |  | 0.57 |
|  | Vermis Crus I | 1.09 |  | > 0.01 | 0.06 |  | 0.10 |
|  | Vermis Crus II | 3.65 |  |  | 0.03 |  | 0.07 |
|  | Vermis VIIb | 0.18 |  | > 0.01 |  |  | > 0.01 |
|  | Vermis VIIIa | 0.05 |  | 0.03 |  | 0.01 |  |
|  | Vermis VIIIb | 0.03 | 0.04 | 0.01 |  | 0.10 |  |
|  | Vermis IX | 2.04 | 0.31 |  |  | 2.36 |  |
|  | Vermis X | 0.65 | 0.03 |  |  | 0.51 |  |

**Supplementary Table 3.** One-sample Kolmogorov-Smirnov test for normality distribution of beta-weights (effective connectivity) of all task contrasts. Significant results are in bold.

|  | One-sample Kolmogorov-Smirnov test | | |
| --- | --- | --- | --- |
|  | *D* value | df | Sig. |
| Emotion | **.06** | **671** | **.00** |
| Language | **.06** | **671** | **.00** |
| Motor | **.05** | **671** | **.00** |
| Relational | **.08** | **671** | **.00** |
| Mentalizing | **.07** | **671** | **.00** |
| Working Memory | **.07** | **671** | **.00** |

**Supplementary Table 4.** Spearman’s rho correlations between task contrast activations. Significant results are in bold.

|  | Emotion | Language | Motor | Relational | Mentalizing | Working Memory |
| --- | --- | --- | --- | --- | --- | --- |
| Emotion |  |  |  |  |  |  |
| Language | **.12^**^** |  |  |  |  |  |
| Motor | -.01 | -.05 |  |  |  |  |
| Relational | **.10^**^** | .07 | .07 |  |  |  |
| Mentalizing | .01 | **.14^**^** | .05 | .07 |  |  |
| Working Memory | .05 | .05 | **.15^**^** | **.22^**^** | **.18^**^** |  |

**Correlation is significant at the 0.01 level (two-tailed).

**Supplementary Table 5.** Wilcoxon signed-rank test results for all possible task pairs. Significant results are in bold.

|  | Negative ranks | | |  | Positive ranks | | |  | Test statistics | | |
| --- | --- | --- | --- | --- | --- | --- | --- | --- | --- | --- | --- |
|  | *n* | Mean Rank | Sum of Ranks |  | *n* | Mean Rank | Sum of Ranks |  | Ties | *Z* | *p* |
| (language)–(emotion) | **272** | **317.63** | **86394** |  | **399** | **348.53** | **139062** |  | **0** | **–5.24^b^** | **.00*** |
| (motor)–(emotion) | **83** | **16.72** | **13340** |  | **588** | **36.74** | **212116** |  | **0** | **–19.79^b^** | **.00*** |
| (relational)–(emotion) | 345 | 321.00 | 110745 |  | 326 | 351.87 | 114711 |  | 0 | –.40^b^ | .69 |
| (mentalizing)–(emotion) | **219** | **276.57** | **60569** |  | **452** | **364.79** | **164887** |  | **0** | **–1.38^b^** | **.00*** |
| (wm)–(emotion) | **67** | **114.73** | **7687** |  | **604** | **36.54** | **217769** |  | **0** | **–2.91^b^** | **.00*** |
| (motor)–(language) | **115** | **191.50** | **22022** |  | **556** | **365.89** | **203434** |  | **0** | **–18.06^b^** | **.00*** |
| (relational)–(language) | **381** | **342.47** | **130482.50** |  | **290** | **327.49** | **94973.50** |  | **0** | **–3.54^a^** | **.00*** |
| (mentalizing)–(language) | **264** | **297.78** | **78613** |  | **407** | **36.79** | **146843** |  | **0** | **–6.79^a^** | **.00*** |
| (wm)–(language) | **82** | **15.80** | **12366** |  | **589** | **361.78** | **213090** |  | **0** | **–19.98^a^** | **.00*** |
| (relational)–(motor) | **577** | **359.66** | **207521** |  | **94** | **19.80** | **17935** |  | **0** | **–18.87^b^** | **.00*** |
| (mentalizing)–(motor) | **513** | **373.46** | **191586** |  | **158** | **214.37** | **33870** |  | **0** | **–15.70^b^** | **.00*** |
| (wm)–(motor) | **250** | **294.76** | **73691** |  | **421** | **36.49** | **151765** |  | **0** | **–7.77^a^** | **.00*** |
| (mentalizing)–(relational) | **222** | **308.44** | **68474** |  | **449** | **349.63** | **156982** |  | **0** | **–8.81^a^** | **.00*** |
| (wm)–(relational) | **69** | **11.12** | **7598** |  | **602** | **361.89** | **217858** |  | **0** | **–2.93^a^** | **.00*** |
| (wm)–(mentalizing) | **104** | **168.42** | **17516** |  | **567** | **366.74** | **207940** |  | **0** | **–18.96^a^** | **.00*** |

*Indicates statistically significant change

^a^Based on negative ranks

^b^Based on positive ranks

**Supplementary Table 6.** Paired samples t-tests on individual beta-weights (effective connectivity) for all proximal ROIs. Significant results are in bold.

|  | Mean | Std Dev | Paired *t* test | | |
| --- | --- | --- | --- | --- | --- |
|  |  |  | *t* value | df | Sig. (two-tailed) |
| *L5 – L5a* |  |  |  |  |  |
| L5_R_ATL | .02 | .49 | -1.18 | 670 | .24 |
| L5a_R_ATL | .04 | .46 |  |  |  |
| L5_R_DMPFC | -.05 | .50 | .44 | 670 | .66 |
| L5a_R_DMPFC | -.06 | .51 |  |  |  |
| L5_R_PreC | .08 | .51 | -1.65 | 670 | .10 |
| L5a_R_PreC | .12 | .57 |  |  |  |
| L5_R_TPJ | .07 | .55 | 1.64 | 670 | .10 |
| L5a_R_TPJ | .03 | .55 |  |  |  |
| L5_R_VMPFC | .04 | .45 | -.44 | 670 | .66 |
| L5a_R_VMPFC | .05 | .47 |  |  |  |
| *L6 – L6a* |  |  |  |  |  |
| L6_R_ATL | .07 | .47 | .89 | 670 | .38 |
| L6a_R_ATL | .05 | .47 |  |  |  |
| L6_R_DMPFC | .03 | .46 | -1.08 | 670 | .28 |
| L6a_R_DMPFC | .06 | .47 |  |  |  |
| **L6_R_PreC** | **.08** | **.51** | **5.72** | **670** | **.00** |
| **L6a_R_PreC** | **-.04** | **.49** |  |  |  |
| **L6_R_VMPFC** | **.01** | **.47** | **-4.23** | **670** | **.00** |
| **L6a_R_VMPFC** | **.10** | **.49** |  |  |  |
| *L7 – L7a* |  |  |  |  |  |
| **L7_R_ATL** | **.09** | **.49** | **4.81** | **670** | **.00** |
| **L7a_R_ATL** | **-.01** | **.55** |  |  |  |
| L7_R_DMPFC | .08 | .51 | 1.71 | 670 | .09 |
| L7a_R_DMPFC | .05 | .50 |  |  |  |
| **L7_R_PreC** | **.03** | **.53** | **7.31** | **670** | **.00** |
| **L7a_R_PreC** | **-.11** | **.56** |  |  |  |
| **L7_R_TPJ** | **.10** | **.57** | **8.81** | **670** | **.00** |
| **L7a_R_TPJ** | **-.07** | **.54** |  |  |  |
| **L7_R_VMPFC** | **.18** | **.54** | **4.77** | **670** | **.00** |
| **L7a_R_VMPFC** | **.09** | **.49** |  |  |  |
| *R1 – R1a* |  |  |  |  |  |
| R1_L_ATL | .07 | .46 | -.72 | 670 | .47 |
| R1a_L_ATL | .09 | .51 |  |  |  |
| R1_L_DMPFC | .02 | .50 | -1.13 | 670 | .26 |
| R1a_L_DMPFC | .05 | .47 |  |  |  |
| R1_L_TPJ | .09 | .51 | -1.14 | 670 | .26 |
| R1a_L_TPJ | .11 | .52 |  |  |  |
| R1_L_VMPFC | .06 | .48 | 1.08 | 670 | .28 |
| R1a_L_VMPFC | .04 | .47 |  |  |  |

**Supplementary Table 7.** Wilcoxon signed ranks test on individual beta-weights (effective connectivity) for proximal ROIs not meeting normality criteria. Significant results are in bold.

|  | Wilcoxon signed ranks test | |
| --- | --- | --- |
|  | *Z* value | Sig. (two-tailed) |
| *L6 – L6a* |  |  |
| L6_R_TPJ | -1.84 | .07 |
| L6a_R_TPJ |  |  |
| *L7 – L7a* |  |  |
| **L7_R_TPJ** | **-8.41** | **.00** |
| **L7a_R_TPJ** |  |  |
| *R1 – R1a* |  |  |
| R1_L_PreC | -1 | .32 |
| R1a_L_PreC |  |  |

**Supplementary Table 8.** Motor mentalizing. FDR-corrected p-values for t-tests for all cerebello-cerebral mentalizing, and cerebellar motor to cerebral mentalizing connections.

|  | Mentalizing | |  | Motor | |  | Mentalizing | |  | Motor | |
| --- | --- | --- | --- | --- | --- | --- | --- | --- | --- | --- | --- |
| Cerebellar  lobes | Effective connections | FDR corrected p-values |  | Effective connections | FDR corrected p-values |  | Effective connections | FDR corrected  p-values |  | Effective connections | FDR corrected  p-values |
| Anterior lobe |  |  |  | L6-R ATL | 0.08 |  |  |  |  | **R6-L ATL** | **0** |
|  |  |  |  | L6-R DMPFC | 0.1723077 |  |  |  |  | R6-L DMPFC | 0.8797727 |
|  |  |  |  | L6-R PreC | 0.1582609 |  |  |  |  | R6-L PreC | 0.3955 |
|  |  |  |  | **L6-R TPJ** | **0.00777778** |  |  |  |  | R6-L TPJ | 0.707 |
|  |  |  |  | L6-R VMPFC | 0.3125862 |  |  |  |  | R6-L VMPFC | 0.4231818 |
| Superior  posterior lobe | **L1-R ATL** | **0** |  | L1-R ATL | 0.1582609 |  | **R1-L ATL** | **0** |  | R1-L ATL | 0.707 |
|  | **L1-R DMPFC** | **0** |  | L1-R DMPFC | 0.4801563 |  | R1-L DMPFC | 0.3123077 |  | **R1-L DMPFC** | **0.028** |
|  | L1-R PreC | 0.187234 |  | **L1-R PreC** | **0** |  | R1-L PreC | 0.1320455 |  | **R1-L PreC** | **0** |
|  | **L1-R TPJ** | **0** |  | **L1-R TPJ** | **0.02545455** |  | **R1-L TPJ** | **0** |  | R1-L TPJ | 0.07 |
|  | **L1-R VMPFC** | **0.01375** |  | L1-R VMPFC | 0.15 |  | **R1-L VMPFC** | **0.0035** |  | R1-L VMPFC | 0.91875 |
|  | **L2-R ATL** | **0** |  | L2-R ATL | 0.14525 |  | **R1a-L ATL** | **0** |  | R2-L ATL | 0.91875 |
|  | **L2-R DMPFC** | **0** |  | L2-R DMPFC | 0.1624 |  | **R1a-L DMPFC** | **0.03** |  | R2-L DMPFC | 0.91875 |
|  | **L2-R PreC** | **0** |  | L2-R PreC | 0.4312903 |  | R1a-L PreC | 0.8017188 |  | R2-L PreC | 0.7566667 |
|  | **L2-R TPJ** | **0** |  | **L2-R TPJ** | **0.0105** |  | **R1a-L TPJ** | **0** |  | R2-L TPJ | 0.336875 |
|  | L2-R VMPFC | 0.09044444 |  | L2-R VMPFC | 0.1326316 |  | **R1a-L VMPFC** | **0.04433333** |  | **R2-L VMPFC** | **0.02625** |
|  | **L3-R ATL** | **0** |  | L5-R ATL | 0.588 |  | R2-L ATL | 0.736129 |  | R5-L ATL | 0.994 |
|  | **L3-R DMPFC** | **0** |  | L5-R DMPFC | 0.1326316 |  | R2-L DMPFC | 0.06125 |  | **R5-L DMPFC** | **0** |
|  | **L3-R PreC** | **0.01132353** |  | L5-R PreC | 0.126 |  | R2-L PreC | 0.6591667 |  | R5-L PreC | 0.4469231 |
|  | **L3-R TPJ** | **0** |  | **L5-R TPJ** | **0** |  | **R2-L TPJ** | **0.02961538** |  | R5-L TPJ | 0.336875 |
|  | **L3-R VMPFC** | **0** |  | **L5-R VMPFC** | **0** |  | R2-L VMPFC | 0.5068966 |  | R5-L VMPFC | 0.91875 |
|  | **L4-R ATL** | **0** |  |  |  |  | R3-L ATL | 0.07368421 |  | R7-L ATL | 0.3955 |
|  | **L4-R DMPFC** | **0.03025** |  |  |  |  | R3-L DMPFC | 0.3538889 |  | R7-L DMPFC | 0.5506667 |
|  | L4-R PreC | 0.4141176 |  |  |  |  | **R3-L PreC** | **0** |  | R7-L PreC | 0.4469231 |
|  | **L4-R TPJ** | **0.006875** |  |  |  |  | **R3-L TPJ** | **0** |  | R7-L TPJ | 0.994 |
|  | **L4-R VMPFC** | **0.04583333** |  |  |  |  | R3-L VMPFC | 0.48875 |  | R7-L VMPFC | 0.91875 |
|  | **L7-R ATL** | **0** |  |  |  |  | **R4-L ATL** | **0** |  |  |  |
|  | **L7-R DMPFC** | **0** |  |  |  |  | R4-L DMPFC | 0.07368421 |  |  |  |
|  | L7-R PreC | 0.2383333 |  |  |  |  | R4-L PreC | 0.951 |  |  |  |
|  | **L7-R TPJ** | **0** |  |  |  |  | R4-L TPJ | 0.1954167 |  |  |  |
|  | **L7-R VMPFC** | **0** |  |  |  |  | **R4-L VMPFC** | **0** |  |  |  |
|  | L7a-R ATL | 0.741 |  |  |  |  |  |  |  |  |  |
|  | **L7a-R DMPFC** | **0.01635135** |  |  |  |  |  |  |  |  |  |
|  | **L7a-R PreC** | **0** |  |  |  |  |  |  |  |  |  |
|  | **L7a-R TPJ** | **0.002115385** |  |  |  |  |  |  |  |  |  |
|  | **L7a-R VMPFC** | **0** |  |  |  |  |  |  |  |  |  |
| Inferior  posterior lobe | L5-R ATL | 0.4347115 |  | **L3-R ATL** | **0** |  | **R5-L ATL** | **0.01272727** |  | R3-L ATL | 0.9566667 |
|  | **L5-R DMPFC** | **0.02315789** |  | L3-R DMPFC | 0.08 |  | R5-L DMPFC | 0.0805 |  | R3-L DMPFC | 0.91875 |
|  | **L5-R PreC** | **0** |  | L3-R PreC | 0.29625 |  | R5-L PreC | 0.06382353 |  | R3-L PreC | 0.5506667 |
|  | **L5-R TPJ** | **0.002115385** |  | L3-R TPJ | 0.1290625 |  | **R5-L TPJ** | **0** |  | R3-L TPJ | 0.91875 |
|  | **L5-R VMPFC** | **0.03219512** |  | L3-R VMPFC | 0.1589583 |  | R5-L VMPFC | 0.3123077 |  | R3-L VMPFC | 0.91875 |
|  | **L5a-R ATL** | **0.02820513** |  | **L3a-R ATL** | **0** |  |  |  |  | R4-L ATL | 0.6444118 |
|  | **L5a-R DMPFC** | **0.008333333** |  | L3a-R DMPFC | 0.05833333 |  |  |  |  | R4-L DMPFC | 0.91875 |
|  | **L5a-R PreC** | **0** |  | **L3a-R PreC** | **0** |  |  |  |  | R4-L PreC | 0.6365625 |
|  | L5a-R TPJ | 0.1733696 |  | **L3a-R TPJ** | **0** |  |  |  |  | R4-L TPJ | 0.707 |
|  | **L5a-R VMPFC** | **0.01257143** |  | L3a-R VMPFC | 0.5661765 |  |  |  |  | R4-L VMPFC | 0.91875 |
|  | **L8-R ATL** | **0.003666667** |  | L4-R ATL | 0.519697 |  |  |  |  |  |  |
|  | **L8-R DMPFC** | **0.003666667** |  | **L4-R DMPFC** | **0** |  |  |  |  |  |  |
|  | L8-R PreC | 0.741 |  | L4-R PreC | 0.4305 |  |  |  |  |  |  |
|  | L8-R TPJ | 0.4029592 |  | L4-R TPJ | 0.1317647 |  |  |  |  |  |  |
|  | L8-R VMPFC | 0.4141176 |  | L4-R VMPFC | 0.2294444 |  |  |  |  |  |  |
| Flocculonodular  lobe | **L6-R ATL** | **0** |  |  |  |  | R6-L ATL | 0.1320455 |  |  |  |
|  | L6-R DMPFC | 0.0625 |  |  |  |  | R6-L DMPFC | 0.1643478 |  |  |  |
|  | **L6-R PreC** | **0** |  |  |  |  | R6-L PreC | 0.8251515 |  |  |  |
|  | **L6-R TPJ** | **0.003666667** |  |  |  |  | **R6-L TPJ** | **0.0175** |  |  |  |
|  | L6-R VMPFC | 0.455566 |  |  |  |  | R6-L VMPFC | 0.8338235 |  |  |  |
|  | **L6a-R ATL** | **0.006875** |  |  |  |  |  |  |  |  |  |
|  | **L6a-R DMPFC** | **0.003666667** |  |  |  |  |  |  |  |  |  |
|  | L6a-R PreC | 0.055 |  |  |  |  |  |  |  |  |  |
|  | **L6a-R TPJ** | **0** |  |  |  |  |  |  |  |  |  |
|  | **L6a-R VMPFC** | **0** |  |  |  |  |  |  |  |  |  |

**Supplementary Table 9.** Cerebello-thalamo-cortical white matter pathways average streamline counts. Results for each cerebellar lobe and the averaged whole CTC and CPC are in bold.

|  | Streamline count | |  | Streamline count | |
| --- | --- | --- | --- | --- | --- |
|  | Mean | SD |  | Mean | SD |
| **Left CTC** | **35.52** | **(13.67)** | **Right CTC** | **32.76** | **(35.73)** |
| **Left superior posterior lobe** | **24.27** | **(26.93)** | **Right superior posterior lobe** | **32.26** | **(35.79)** |
| L1-R_ATL | 7.26 | (13.67) | R1-L_ATL | 8.75 | (21.19) |
| L1-R_DMPFC | 80.17 | (84.47) | R1-L_DMPFC | 100.40 | (98.79) |
| L1-R_PreC | 21.18 | (36.19) | R1-L_PreC | 29.22 | (38.89) |
| L1-R_TPJ | 4.65 | (10.62) | R1-L_TPJ | 4.85 | (9.99) |
| L1-R_VMPFC | 33.47 | (47.37) | R1-L_VMPFC | 38.30 | (66.03) |
|  |  |  | R1a-L_ATL | 8.86 | (21.61) |
|  |  |  | R1a-L_DMPFC | 98.88 | (94.63) |
|  |  |  | R1a-L_PreC | 29.43 | (39.13) |
|  |  |  | R1a-L_TPJ | 4.52 | (9.72) |
|  |  |  | R1a-L_VMPFC | 39.21 | (68.69) |
| L2-R_ATL | 9.62 | (17.18) | R2-L_ATL | 11.65 | (21.67) |
| L2-R_DMPFC | 112.91 | (128.06) | R2-L_DMPFC | 143.06 | (160.77) |
| L2-R_PreC | 28.25 | (40.34) | R2-L_PreC | 41.97 | (51.97) |
| L2-R_TPJ | 7.22 | (20.47) | R2-L_TPJ | 6.96 | (13.21) |
| L2-R_VMPFC | 48.28 | (77.53) | R2-L_VMPFC | 54.87 | (90.42) |
| L3-R_ATL | 4.01 | (8.18) | R3-L_ATL | 3.62 | (8.80) |
| L3-R_DMPFC | 48.55 | (65.79) | R3-L_DMPFC | 42.08 | (50.82) |
| L3-R_PreC | 12.39 | (22.01) | R3-L_PreC | 13.06 | (17.96) |
| L3-R_TPJ | 2.73 | (6.27) | R3-L_TPJ | 2.03 | (4.71) |
| L3-R_VMPFC | 21.2 | (45.41) | R3-L_VMPFC | 16.25 | (26.35) |
| L4-R_ATL | 6.81 | (10.49) | R4-L_ATL | 5.23 | (10.85) |
| L4-R_DMPFC | 78.37 | (90.26) | R4-L_DMPFC | 58.33 | (69.50) |
| L4-R_PreC | 21.13 | (41.09) | R4-L_PreC | 18.83 | (31.59) |
| L4-R_TPJ | 4.82 | (12.42) | R4-L_TPJ | 2.84 | (6.52) |
| L4-R_VMPFC | 33.49 | (53.01) | R4-L_VMPFC | 23.23 | (43.42) |
| L7-R_ATL | 3.23 | (6.86) |  |  |  |
| L7-R_DMPFC | 39.76 | (60.18) |  |  |  |
| L7-R_PreC | 9.66 | (16.15) |  |  |  |
| L7-R_TPJ | 2.25 | (5.72) |  |  |  |
| L7-R_VMPFC | 16.47 | (31.70) |  |  |  |
| L7a-R_ATL | 3.1 | (6.40) |  |  |  |
| L7a-R_DMPFC | 39.03 | (56.93) |  |  |  |
| L7a-R_PreC | 9.92 | (18.13) |  |  |  |
| L7a-R_TPJ | 2.24 | (5.78) |  |  |  |
| L7a-R_VMPFC | 15.87 | (30.79) |  |  |  |
| **Left inferior posterior lobe** | **38.42** | **(41.90)** | **Right inferior posterior lobe** | **22.04** | **(22.76)** |
| L5-R_ATL | 6.25 | (10.14) | R5-L_ATL | 5.42 | (11.28) |
| L5-R_DMPFC | 71 | (80.34) | R5-L_DMPFC | 59.57 | (70.72) |
| L5-R_PreC | 19.27 | (29.65) | R5-L_PreC | 17.96 | (24.36) |
| L5-R_TPJ | 4.23 | (10.39) | R5-L_TPJ | 2.91 | (6.33) |
| L5-R_VMPFC | 31.01 | (52.30) | R5-L_VMPFC | 24.34 | (44.99) |
| L5a-R_ATL | 11.78 | (17.70) |  |  |  |
| L5a-R_DMPFC | 139.34 | (134.45) |  |  |  |
| L5a-R_PreC | 34.61 | (55.27) |  |  |  |
| L5a-R_TPJ | 9.32 | (23.67) |  |  |  |
| L5a-R_VMPFC | 57.38 | (79.88) |  |  |  |
| **Left flocculonodular lobe** | **56.10** | **(58.60)** | **Right flocculonodular lobe** | **45.99** | **(47.81)** |
| L6-R_ATL | 11.46 | (15.54) | R6-L_ATL | 11.31 | (22.76) |
| L6-R_DMPFC | 132.81 | (125.52) | R6-L_DMPFC | 125.54 | (110.95) |
| L6-R_PreC | 35.59 | (50.65) | R6-L_PreC | 39.66 | (49.49) |
| L6-R_TPJ | 8.86 | (24.91) | R6-L_TPJ | 6.37 | (10.73) |
| L6-R_VMPFC | 55.38 | (68.56) | R6-L_VMPFC | 47.10 | (58.77) |
| L6a-R_ATL | 10.06 | (13.35) |  |  |  |
| L6a-R_DMPFC | 117.07 | (109.76) |  |  |  |
| L6a-R_PreC | 31.78 | (48.23) |  |  |  |
| L6a-R_TPJ | 8.11 | (27.19) |  |  |  |
| L6a-R_VMPFC | 49.1 | (60.73) |  |  |  |
| L7-R_ATL | 16.99 | (25.48) |  |  |  |
| L7-R_DMPFC | 211.01 | (179.70) |  |  |  |
| L7-R_PreC | 50.41 | (59.08) |  |  |  |
| L7-R_TPJ | 11.72 | (27.44) |  |  |  |
| L7-R_VMPFC | 91.22 | (113.75) |  |  |  |

**Supplementary Table 10.** Cortico-ponto-cerebellar white matter pathways average streamline counts. Results for each cerebellar lobe and the averaged whole CTC and CPC are in bold.

|  | Streamline count | |  | Streamline count | |
| --- | --- | --- | --- | --- | --- |
|  | Mean | SD |  | Mean | SD |
| **Left CPC** | **28.04** | **(33.81)** | **Right CPC** | **21.74** | **(25.30)** |
| **Left superior posterior lobe** | **17.82** | **(19.26)** | **Right superior posterior lobe** | **20.89** | **(19.93)** |
| R_ATL-L1 | 15.96 | (17.86) | L_ATL-R1 | 12.99 | (15.50) |
| R_DMPFC-L1 | 72.64 | (98.81) | L_DMPFC-R1 | 72.82 | (87.45) |
| R_PreC-L1 | 10.92 | (13.11) | L_PreC-R1 | 19.85 | (21.81) |
| R_TPJ-L1 | 23.08 | (33.07) | L_TPJ-R1 | 15.34 | (29.55) |
| R_VMPFC-L1 | 21.38 | (42.25) | L_VMPFC-R1 | 18.76 | (38.54) |
|  |  |  | L_ATL-R1a | 12.32 | (13.39) |
|  |  |  | L_DMPFC-R1a | 67.38 | (78.16) |
|  |  |  | L_PreC-R1a | 18.78 | (21.52) |
|  |  |  | L_TPJ-R1a | 14.73 | (26.95) |
|  |  |  | L_VMPFC-R1a | 18.50 | (39.59) |
| R_ATL-L2 | 15.81 | (17.84) | L_ATL-R2 | 11.79 | (13.64) |
| R_DMPFC-L2 | 71.99 | (84.57) | L_DMPFC-R2 | 66.68 | (87.66) |
| R_PreC-L2 | 13.32 | (27.04) | L_PreC-R2 | 19.02 | (24.39) |
| R_TPJ-L2 | 23.77 | (39.17) | L_TPJ-R2 | 13.84 | (20.04) |
| R_VMPFC-L2 | 18.80 | (33.76) | L_VMPFC-R2 | 15.96 | (34.09) |
| R_ATL-L3 | 5.37 | (7.12) | L_ATL-R3 | 3.42 | (5.37) |
| R_DMPFC-L3 | 25.25 | (40.25) | L_DMPFC-R3 | 20.73 | (30.74) |
| R_PreC-L3 | 4.33 | (7.97) | L_PreC-R3 | 5.26 | (8.02) |
| R_TPJ-L3 | 8.23 | (15.01) | L_TPJ-R3 | 4.21 | (9.36) |
| R_VMPFC-L3 | 7.06 | (26.53) | L_VMPFC-R3 | 4.02 | (9.31) |
| R_ATL-L4 | 15.66 | (18.74) | L_ATL-R4 | 7.92 | (10.16) |
| R_DMPFC-L4 | 66.82 | (89.37) | L_DMPFC-R4 | 45.59 | (57.27) |
| R_PreC-L4 | 10.35 | (13.65) | L_PreC-R4 | 12.09 | (14.93) |
| R_TPJ-L4 | 24.00 | (35.34) | L_TPJ-R4 | 9.73 | (23.76) |
| R_VMPFC-L4 | 17.47 | (36.53) | L_VMPFC-R4 | 10.48 | (22.34) |
| R_ATL-L7 | 4.26 | (6.60) |  |  |  |
| R_DMPFC-L7 | 17.58 | (26.19) |  |  |  |
| R_PreC-L7 | 3.16 | (5.54) |  |  |  |
| R_TPJ-L7 | 6.36 | (11.97) |  |  |  |
| R_VMPFC-L7 | 4.62 | (11.40) |  |  |  |
| R_ATL-L7a | 3.25 | (5.17) |  |  |  |
| R_DMPFC-L7a | 12.98 | (19.31) |  |  |  |
| R_PreC-L7a | 2.30 | (4.18) |  |  |  |
| R_TPJ-L7a | 4.58 | (7.69) |  |  |  |
| R_VMPFC-L7a | 3.38 | (10.53) |  |  |  |
| **Left inferior posterior lobe** | **13.96** | **(14.36)** | **Right inferior posterior lobe** | **1.86** | **(1.62)** |
| R_ATL-L5 | 4.15 | (5.85) | L_ATL-R5 | 0.93 | (2.51) |
| R_DMPFC-L5 | 17.68 | (24.50) | L_DMPFC-R5 | 4.75 | (7.81) |
| R_PreC-L5 | 2.82 | (4.14) | L_PreC-R5 | 1.19 | (2.18) |
| R_TPJ-L5 | 5.89 | (8.96) | L_TPJ-R5 | 1.05 | (3.25) |
| R_VMPFC-L5 | 6.52 | (16.59) | L_VMPFC-R5 | 1.40 | (4.03) |
| R_ATL-L5a | 10.20 | (11.47) |  |  |  |
| R_DMPFC-L5a | 51.66 | (68.38) |  |  |  |
| R_PreC-L5a | 7.57 | (9.46) |  |  |  |
| R_TPJ-L5a | 14.23 | (17.50) |  |  |  |
| R_VMPFC-L5a | 18.91 | (33.61) |  |  |  |
| **Left flocculonodular lobe** | **57.84** | **(46.81)** | **Right flocculonodular lobe** | **45.87** | **(42.73)** |
| R_ATL-L6 | 34.74 | (36.06) | L_ATL-R6 | 21.82 | (22.65) |
| R_DMPFC-L6 | 138.17 | (170.01) | L_DMPFC-R6 | 121.89 | (140.21) |
| R_PreC-L6 | 22.50 | (26.27) | L_PreC-R6 | 33.73 | (34.68) |
| R_TPJ-L6 | 54.44 | (72.23) | L_TPJ-R6 | 27.97 | (69.82) |
| R_VMPFC-L6 | 33.34 | (48.08) | L_VMPFC-R6 | 23.94 | (42.36) |
| R_ATL-L6a | 38.34 | (43.20) |  |  |  |
| R_DMPFC-L6a | 144.61 | (164.80) |  |  |  |
| R_PreC-L6a | 25.14 | (30.59) |  |  |  |
| R_TPJ-L6a | 60.58 | (81.73) |  |  |  |
| R_VMPFC-L6a | 31.18 | (44.83) |  |  |  |
| R_ATL-L7 | 22.69 | (6.60) |  |  |  |
| R_DMPFC-L7 | 150.01 | (26.19) |  |  |  |
| R_PreC-L7 | 20.27 | (5.54) |  |  |  |
| R_TPJ-L7 | 27.15 | (11.97) |  |  |  |
| R_VMPFC-L7 | 64.50 | (11.40) |  |  |  |

**Supplementary Table 11.** One-sample Kolmogorov-Smirnov test for normality distribution of cerebello-thalamo-cortical and cortico-ponto-cerebellar white matter pathway streamline counts (cerebellar lobe averages). Significant results are in bold.

|  |  | One-sample Kolmogorov-Smirnov test | | |
| --- | --- | --- | --- | --- |
|  |  | *D* value | df | Sig. |
| CTC | Left cerebellum to right cerebrum | **0.12** | **671** | **.00** |
|  | Superior posterior lobe | **0.16** | **671** | **.00** |
|  | Inferior posterior lobe | **0.14** | **671** | **.00** |
|  | Flocculonodular lobe | **0.12** | **671** | **.00** |
|  | Right cerebellum to left cerebrum | **0.14** | **671** | **.00** |
|  | Superior posterior lobe | **0.14** | **671** | **.00** |
|  | Inferior posterior lobe | **0.19** | **671** | **.00** |
|  | Flocculonodular lobe | **0.13** | **671** | **.00** |
| CPC | Left cerebrum to right cerebellum | **0.14** | **671** | **.00** |
|  | Superior posterior lobe | **0.14** | **671** | **.00** |
|  | Inferior posterior lobe | **0.25** | **671** | **.00** |
|  | Flocculonodular lobe | **0.15** | **671** | **.00** |
|  | Right cerebrum to left cerebellum | **0.14** | **671** | **.00** |
|  | Superior posterior lobe | **0.14** | **671** | **.00** |
|  | Inferior posterior lobe | **0.17** | **671** | **.00** |
|  | Flocculonodular lobe | **0.14** | **671** | **.00** |

**Supplementary Table 12.** Spearman’s rho correlations between cerebello-thalamo-cortical streamline counts averaged across cerebellar lobes. Bold indicates the correlation of interest (e.g. left superior lobe correlated to right superior lobe streamline count). Black boxes enclose correlations between right-left CTC streamline counts.

|  |  | Left cerebellum to right cerebrum | | |  | Right cerebellum to left cerebrum | | |
| --- | --- | --- | --- | --- | --- | --- | --- | --- |
|  |  | Superior posterior lobe | Inferior posterior lobe | Flocculo  nodular  lobe |  | Superior posterior lobe | Inferior posterior lobe | Flocculo  nodular  lobe |
| Left cerebellum to right cerebrum | Superior posterior lobe | 1 | .642^**^ | .774^**^ |  | **.230^**^** | .121^**^ | .191^**^ |
|  | Inferior posterior lobe | .642^**^ | 1 | .845^**^ |  | .148^**^ | **.275^**^** | .179^**^ |
|  | Flocculonodular lobe | .774^**^ | .845^**^ | 1 |  | .179^**^ | .162^**^ | **.234^**^** |
| Right cerebellum to left cerebrum | Superior posterior lobe | **.230^**^** | .148^**^ | .179^**^ |  | 1 | .534^**^ | .697^**^ |
|  | Inferior posterior lobe | .121^**^ | **.275^**^** | .162^**^ |  | .534^**^ | 1 | .599^**^ |
|  | Flocculonodular lobe | .191^**^ | .179^**^ | **.234^**^** |  | .697^**^ | .599^**^ | 1 |

**Correlation is significant at the 0.01 level (two-tailed).

**Supplementary Table 13.** Spearman’s rho correlations between cortico-ponto-cerebellar streamline counts averaged across cerebellar lobes. Bold indicates the correlation of interest (eg. left superior lobe correlated to right superior lobe streamline count). Black boxes enclose correlations between right-left CPC streamline counts.

|  |  | Left cerebrum to right cerebellum | | |  | Right cerebrum to left cerebellum | | |
| --- | --- | --- | --- | --- | --- | --- | --- | --- |
|  |  | Superior posterior lobe | Inferior posterior lobe | Flocculo  nodular lobe |  | Superior posterior lobe | Inferior posterior lobe | Flocculo  nodular lobe |
| Left cerebrum to right cerebellum | Superior posterior lobe | 1 | .557^**^ | .825^**^ |  | **.240^**^** | .201^**^ | .204^**^ |
|  | Inferior posterior lobe | .557^**^ | 1 | .641^**^ |  | .106^**^ | **.251^**^** | .160^**^ |
|  | Flocculonodular lobe | .825^**^ | .641^**^ | 1 |  | .164^**^ | .207^**^ | **.215^**^** |
| Right cerebrum to left cerebellum | Superior posterior lobe | **.240^**^** | .106^**^ | .164^**^ |  | 1 | .688^**^ | .860^**^ |
|  | Inferior posterior lobe | .201^**^ | **.251^**^** | .207^**^ |  | .688^**^ | 1 | .829^**^ |
|  | Flocculonodular lobe | .204^**^ | .160^**^ | **.215^**^** |  | .860^**^ | .829^**^ | 1 |

**Correlation is significant at the 0.01 level (two-tailed).

**Supplementary Table 14**. Wilcoxon signed-rank test results for all possible left-right streamline count cerebellar hemisphere and lobe pairs. Significant results are in bold.

|  | Negative ranks | | |  | Positive ranks | | |  | Test statistics | | |
| --- | --- | --- | --- | --- | --- | --- | --- | --- | --- | --- | --- |
|  | *n* | Mean Rank | Sum of Ranks |  | *n* | Mean Rank | Sum of Ranks |  | Ties | *Z* | *p* |
| CTC R – CTC L | **372** | **340.75** | **126759.50** |  | **297** | **327.80** | **97355.50** |  | **2** | **–2.94^b^** | **.003*** |
| (SPR)–(SPL) | **235** | **297.57** | **69930** |  | **433** | **354.54** | **153516** |  | **3** | **–8.38^a^** | **.00*** |
| (IPR)–(IPL) | **503** | **359.68** | **180919** |  | **166** | **260.22** | **43196** |  | **2** | **–13.77^b^** | **.00*** |
| (FR)–(FL) | **429** | **341.68** | **146581** |  | **240** | **323.06** | **77534** |  | **2** | **– 6.90^b^** | **.00*** |
| CPC R – CPC L | **424** | **349.52** | **148197.50** |  | **245** | **309.87** | **75917.50** |  | **2** | **–7.23^b^** | **.00*** |
| (SPR)–(SPL) | **300** | **311.49** | **93446** |  | **369** | **354.12** | **130669** |  | **2** | **–3.72^a^** | **.00*** |
| (IPR)–(IPL) | **649** | **339.66** | **220437.50** |  | **18** | **130.03** | **2340.50** |  | **4** | **–21.91^b^** | **.00*** |
| (FR)–(FL) | **404** | **355.75** | **143724.50** |  | **265** | **303.36** | **80390.50** |  | **2** | **–6.33^b^** | **.00*** |
| CPC R – CTC R | **484** | **357.61** | **173081.50** |  | **184** | **273.72** | **50364.5** |  | **3** | **–12.30^b^** | **.00*** |
| CPC L – CTC L | **434** | **353.97** | **153.624** |  | **235** | **299.96** | **70491** |  | **2** | **–8.31^b^** | **.00*** |

*Indicates statistically significant change

^a^Based on negative ranks

^b^Based on positive ranks

CTC: cerebello-thalamo-cortical pathway, CPC: cortico-ponto-cerebellar pathway, SPR: superior posterior right lobe, SPL: superior posterior left lobe, IPR: inferior posterior right lobe, IPL: inferior posterior left lobe, FR: flocculonodular right lobe, FL: flocculonodular left lobe

**Supplementary Table 15.** Motor control white matter pathways average streamline counts. Results for the averaged whole CTC and CPC are in bold.

|  | Streamline count | |  | Streamline count | |
| --- | --- | --- | --- | --- | --- |
|  | Mean | SD |  | Mean | SD |
| **Left CTC** | **14.51** | **(10.72)** | **Right CTC** | **14.20** | **(10.79)** |
| Left superior posterior lobe | 13.24 | (12.12) | Right superior posterior lobe | 15.88 | (12.27) |
| Left inferior posterior lobe | 16.12 | (14.13) | Right inferior posterior lobe | 15.11 | (16.22) |
| **Left CPC** | **21.07** | **(16.13)** | **Right CPC** | **13.92** | **(12.52)** |
| Left superior posterior lobe | 6.96 | (6.14) | Right superior posterior lobe | 9.90 | (8.18) |
| Left inferior posterior lobe | 12.65 | (14.97) | Right inferior posterior lobe | 10.52 | (11.93) |

**Supplementary Table 16.** Wilcoxon signed-rank test results for all possible motor-mentalizing CTC and CPC streamline count cerebellar lobe pairs. Significant results are in bold.

|  | Negative ranks | | |  | Positive ranks | | |  | Test statistics | | |
| --- | --- | --- | --- | --- | --- | --- | --- | --- | --- | --- | --- |
|  | *n* | Mean Rank | Sum of Ranks |  | *n* | Mean Rank | Sum of Ranks |  | Ties | *Z* | *p* |
| **CTC L m – CTC L M** | **663** | **337.35** | **223661** |  | **6** | **75.67** | **454** |  | **2** | **–22.32^b^** | **.00*** |
| **(SPL m)–(SPL M)** | **617** | **346.48** | **213780.50** |  | **52** | **198.74** | **10334.50** |  | **2** | **–20.34^b^** | **.00*** |
| **(IPL m )–(IPL M)** | **615** | **352.49** | **216783** |  | **54** | **135.78** | **7332** |  | **2** | **–20.94^b^** | **.00*** |
| **CTC R m – CTC L M** | **659** | **339.35** | **223634** |  | **10** | **48.10** | **481** |  | **2** | **–22.31^b^** | **.00*** |
| **(SPR m)–(SPL M)** | **630** | **348.37** | **219474.50** |  | **39** | **118.99** | **4640.50** |  | **2** | **–21.48^b^** | **.00*** |
| **(IPR m)–(IPR M)** | **427** | **355.67** | **151870.50** |  | **235** | **287.59** | **67582.50** |  | **9** | **–8.56^b^** | **.00*** |
| **CPC L m – CPC L M** | **522** | **358.36** | **187064** |  | **147** | **252.05** | **37051** |  | **2** | **–15.00 ^b^** | **.00*** |
| **(SPL m)–(SPL M)** | **666** | **335.86** | **223685** |  | **3** | **143.33** | **430** |  | **2** | **–22.32^b^** | **.00*** |
| **(IPL m)–(IPL M)** | **390** | **342.26** | **133479.50** |  | **278** | **323.62** | **89966.50** |  | **3** | **–4.36^b^** | **.00*** |
| **CPC R m – CPC L M** | **630** | **345.95** | **217945.50** |  | **38** | **144.75** | **5500.50** |  | **3** | **–21.29^b^** | **.00*** |
| **(SPR m)–(SPR M)** | **651** | **343.34** | **223512.50** |  | **18** | **33.47** | **602.50** |  | **2** | **–22.29^b^** | **.00*** |
| **(IPR m)–(IPR M)** | **23** | **128.83** | **2986** |  | **643** | **340.79** | **219125** |  | **5** | **–21.76^a^** | **.00*** |

*Indicates statistically significant change

^a^Based on negative ranks

^b^Based on positive ranks

CTC: cerebello-thalamo-cortical pathway, CPC: cortico-ponto-cerebellar pathway, m: motor, ML mentalizing, SPR: superior posterior right lobe, SPL: superior posterior left lobe, IPR: inferior posterior right lobe, IPL: inferior posterior left lobe, FR: flocculonodular right lobe, FL: flocculonodular left lobe
